# Sweet and sour synergy: exploring the antibacterial and antibiofilm activity of acetic acid and vinegar combined with medical-grade honeys

**DOI:** 10.1101/2023.04.03.535340

**Authors:** Freya Harrison, Anisa Blower, Chris de Wolf, Erin Connelly

**Affiliations:** School of Life Sciences, University of Warwick, Coventry CV4 7AL, United Kingdom; Warwick Integrative Synthetic Biology Centre, University of Warwick, Coventry CV4 7AL, United Kingdom

**Keywords:** Antimicrobials, biofilm, natural products, synergy

## Abstract

*Oxymel*, a combination of honey and vinegar, has been used as a remedy for wounds and infections from antiquity to the present day. While honey is now clinically used to treat infected wounds, this use of a complex, raw natural product (NP) mixture is unusual in modern western medicine. Research into the antimicrobial activity of NPs more usually focusses on finding a single active compound. The acetic acid in vinegar is known to have antibacterial activity at low concentrations and is in clinical use to treat burn wound infections. Here, we investigated the potential for synergistic activity of different compounds present in a complex ingredient used in historical medicine (vinegar) and in an ingredient mixture (*oxymel*). We conducted a systematic review to investigate published evidence for antimicrobial effects of vinegars against human pathogenic bacteria and fungi. No published studies explicitly compared the activity of vinegar with that of a comparable concentration of acetic acid. We then characterised selected vinegars by high-performance liquid chromatography (HPLC) and assessed the antibacterial and antibiofilm activity of the vinegars and acetic acid, alone and in combination with medical-grade honeys, against *P. aeruginosa* and *S. aureus*. We found that some vinegars have antibacterial activity that exceeds that predicted by their acetic acid content alone, but that this depends on the bacterial species being investigated and the growth conditions (media type, planktonic vs. biofilm). Pomegranate vinegars may be particularly interesting candidates for further study. We also conclude that there is potential for acetic acid, and some vinegars, to show synergistic antibiofilm activity with manuka honey.

**Data Summary:** Raw quantitative data and R code for analyses are provided in the supplementary data (Document S1).

## Introduction

Vinegar and honey are ubiquitous ingredients in historical and traditional medical remedies across time and cultures up to the present day (1-5). One influential medieval text describes honey as having ‘the virtues of both food and medicine,’ and, citing Hippocrates and Ibn-Sīnā, the ability to ‘expel, cleanse, nourish and penetrate’ infection, inflammation and wasting conditions when mixed with vinegar in a combination known as *oxymel* or with the addition of water (*hydromel*) (6). This medieval example of *oxymel* as a foundational mixture for remedies and association with ‘cleaning and healing’ of inflamed or infected tissue is indebted to a much earlier medical discourse ranging from Ancient Egypt to the authorities of Antiquity (Hippocrates, Galen, Dioscorides) to the early medieval scholars of Arabic medicine (Ibn-Sīnā) and similar textual and traditional occurrences along the global timeline too numerous to list. In our survey of 40 medieval manuscript sources of 418 Latin, Middle English, Old English, and Welsh recipes from the ninth to the fifteenth centuries, for wounds, skin infections, eye, mouth and throat infections, and all other external infections, there were 151 mentions of vinegar, honey, *oxymel*, or *hydromel*. In contrast, another common historical ingredient of current interest, garlic, only appears 9 times in this dataset (Erin Connelly, manuscript in preparation). Many of these historical uses appear reasonable given current scientific understanding of the antimicrobial activities of both honey and acetic acid – a key component of vinegar – and the wound healing potential of honey.

The acetic acid in vinegar is known to have antibacterial activity at low concentrations, including the ability to kill Gram-positive and Gram-negative opportunistic pathogens living as monospecies biofilms (7-13), and a clinical trial is in progress in the United Kingdom to assess efficacy and optimal dose of acetic acid as a topical treatment for colonised burn wounds (14, 15). A United Kingdom National Survey of burns units performed in October 2019 found that approximately one-third of units used acetic acid-soaked dressings to treat burns infected with *Pseduomonas aeruginosa* and that a daily dressing of 2.5–3% acetic acid is a well-tolerated treatment in these patients (16). Given the presence of many other compounds in vinegar aside from acetic acid, it is possible that some or all vinegars may contain compounds that could enhance the antimicrobial activity of acetic acid by potentiating its effects on bacteria, or by contributing additional antimicrobial activity of their own. While modern drug discovery research that focusses on natural products usually attempts to isolate a single active compound for development into a medicine, it is increasingly recognised that many antimicrobial plant extracts and plant-derived products may owe their activity to combinations of compounds present within them (17). The antimicrobial activity of wine, for instance, which like vinegar contains a range of organic acids, phenolics and alcohol, has been attributed to the combined activity of these different types of compound (13, 18, 19). Although some vinegars with an acetic acid content of only 0.1% can inhibit the growth of a range of bacterial species (20) (7) (12), there is some evidence that the presence of phenolic compounds in fermented vinegars enhance its antimicrobial effect (21). Further research is required to establish the role of polyphenolics and other weak organic acids, such as gallic acid, in contributing to the antimicrobial effect of vinegars, at low concentrations.

Likewise, honey is a complex natural substance that has received extensive research attention for its healing, immunostimulatory and antimicrobial properties. The rationale for the rediscovery of honey by modern medicine has been extensively reviewed (22-24), and numerous trials of honey dressings and gels in wound management have been conducted (25-28). While most attention has focussed on manuka (*Leptospermum*) honey, other honeys have been shown to have good antimicrobial activity (e.g. (29)). Medical honey and honey-impregnated dressings are a common line of care for wound management in clinical settings, including treatment for biofilms, critically colonised wounds and infections with high bacterial levels, chronic ulceration, debridement of dead tissue, malodorous wounds, fungating wounds, burn sites, skin grafts and surgical wounds (30-32).

There are parallel occurrences of honey in historical textual accounts for perilous, non-healing wounds, and even surgical wounds with evidence of use as a preventative measure against post-surgical infections or ulceration. Notably, in many instances honey is combined with vinegar, as the pairwise treatment *oxymel*, or as a compound with medicinal plants, metals, or other ingredients, in ointments, cleansing washes, plasters, and medicated bandages. One Middle English medical recipe collection contains recipes for making *oxymel simple* and *oxymel compound. Oxymel simple* is made of one part honey and two parts vinegar, combined over heat (fire). *Oxymel compound* is *oxymel simple* plus other medicinal ingredients; a common version is *squill oxymel*, which is *oxymel* made with bulbs of the squill, *Drimia maritima* (33). A few examples from select later medieval medical and surgical texts include references to a plaster to prevent the development of an ulcer in a wound (34); to cleanse wounds of dead tissue (35, 36); to remove dead tissue and encourage healing in old *fistulas* (deep-seated ulcer or infection) by applying medicated cloths (*tent*) (37); post-surgical washes after cutting of infected swellings (38); for all manner of wounds in the head and body (39); for *cankers* (wound, ulcer, abscess, sore, swelling) and *festers* (fistula or fistulous wound, abscess, boil) (40-44); mouthwashes for *cankers* in the teeth and gums (45-47); a plaster for an *aposteme* (inflammation, abscess, swelling) (48); and an ointment for itching and pustules in the eyebrows (49).

The broad-spectrum antimicrobial activity of vinegar (7) makes it a promising candidate for further research to test potential synergistic interactions with medical grade honey. Given the lengthy historical tradition, current clinical uses and research outputs for honey and vinegar as single ingredients, and low-cost availability, it is surprising that *oxymel* has not been well studied. Our previous research to detect communities of ingredients in a network of the recipes in one medieval text, showed the existence of a hierarchical structure within the recipes. This resulted in the identification of four core individual ingredients, including honey and vinegar. The results of our pilot experimental data suggest that combinations of honey and vinegar, along with other ingredients, ‘may be worth investigation for their ability to potentiate each other’s antibacterial effects in biofilm models of infection, where the combinatorial activity of agents is more likely to enhance the killing of bacteria with enhanced tolerance’ (50).

We therefore wished to test two hypotheses. First, that all or some vinegars may have antibacterial activity exceeding that of an equivalent concentration of pure acetic acid. Second, that combining either pure acetic acid or vinegar with honey may lead to the identification of a synergistic ability to kill pathogenic bacteria grown as biofilms. We performed a systematic review of the antimicrobial properties of vinegars. We searched for published evidence that included quantitative data on the antimicrobial effect of vinegars against common human bacterial and fungal pathogens and/or against the bacterium *Bacillus subtilis* (because this species is often used as a model Gram-positive organism). Quality screening of articles showed reporting of data to be highly variable in quality. We did not find strong evidence to conclude that the antimicrobial activity of vinegar varies by botanical origin or that the activity of vinegar is entirely due to acetic acid content. We then selected seven types of vinegar, commercially produced from a range of starting materials used to make fermented vinegars, for analysis by reversed-phase high-performance liquid chromatography (HPLC). We then performed an assessment of the antibacterial and antibiofilm activity of these vinegars using *P. aeruginosa* and *S. aureus* as example Gram-negative and Gram-positive bacterial pathogens. Our experimental results show the potential for some vinegars to have antibacterial activity that exceeds that predicted by their acetic acid content alone, but that this depends on the bacterial species being investigated and the growth conditions (media type, planktonic vs. biofilm). Furthermore, there is a potential for acetic acid, and some vinegars, to show synergistic antibiofilm effects with manuka honey. These preliminary results will be enhanced by future testing, by checkerboard analysis for instance, to better understand synergistic implications and applications.

## Materials and methods

### Systematic review

Searches for primary research assessing the antimicrobial nature of vinegar were performed on the 17th – 20th November 2021 using PubMed, Scopus, Web of Science and the Cochrane database of clinical trials. The Boolean search term (“Vinegar” AND (“Antimicrobial” OR “Antibacterial” OR “Antifungal”)) was used to search document title abstract and keyword (for Scopus, Web of Science and Cochrane database of clinical trials). Document title and abstract were searched in PubMed, as it does not allow the targeted searching of keywords. Searches were restricted to exclude reviews (systematic and conference) and editorials, and to include papers of any age or language. Outputs from searches were inputted into one .csv file. Duplicates were then removed manually by using the “sort” function in Microsoft Excel for author, title and DOI.

Abstracts were then read and screened for the presence of primary quantitative data regarding the effects of whole fermented vinegar or acetic acid as an antimicrobial, against human pathogenic bacteria, fungus, and *Bacillus subtilis*. Several papers that investigated “wood vinegar”, which is pyroligneous acid produced by distillation of plant biomass, were excluded at this stage. Further reasons for exclusion at this stage included articles that were non-empirical or inaccessible (due to language or paywall barriers). As only one author assessed each study, 5% of abstracts were re-scored by the same author 4 weeks after the initial screening to assess the consistency of the application of inclusion and exclusion criteria.

Full-text articles of included records were assessed for eligibility. Articles for which the full text was inaccessible due to language, i.e., not written in or translated to English, or paywalled were excluded at this stage (following the request for alternative versions from the authors and translation by our colleagues where possible). Studies that accessed only acetic acid were also excluded. Papers eligible for inclusion were those that contained either minimum inhibitory concentration (MIC) or zone of inhibition (ZOI) data testing the antimicrobial nature of vinegar against human bacterial or fungal pathogens and *Bacillus subtilis*. Data extracted from eligible papers included vinegar type, test microbe (and details of the strain or isolate), assay method and details of the method (growth medium, temperature and agar thickness for ZOI data), reference to diagnostic guidelines, location of data within the article, units, variance/standard deviation, details of positive and negative controls used, and number of replicates conducted. The extracted data was used to select MIC/ZOI data of sufficient quality (containing an appropriate experimental design, negative control, and number of repeats for the technique) for further analysis.

The systematic review identified only one previous paper which considered honey and vinegar as a combination against bacteria (51). The author assayed various honey-vinegar solutions and determined that the combination provides a superior killing effect on select pathogenic bacteria, than vinegar alone. However, because this paper did not include a honey-only treatment, it could not assess whether the combinations of vinegar and honey showed additive or synergistic antibacterial effects. To determine if other papers which assessed combinations of honey and vinegar had been missed by our systematic review, an unrestricted search of PubMed, Cochrane Library, Web of Science, and Scopus for *oxymel* or *oximel* was performed on 13 January 2023. The term *oximel* returned 0 results, while the term *oxymel* revealed a few studies combining *oxymel* with various medicinal plants for obesity-associated inflammation, cardiovascular risk factors, and insulin resistance (52), and the traditional / historical mixture, *squill oxymel* (*Drimia maritima*) for asthma (53), fatty liver (54), and COPD (55), but no antimicrobial investigations.

Two authors repeated the search on 13th (Scopus, PubMed, Cochrane) and 19th (Web of Science) January 2023. The search was date restricted to capture new publications from the search performed in November 2021 to present (January 2023).

### Bacterial Strains and Culture Conditions

*Pseudomonas aeruginosa* PA14 and *Staphylococcus aureus* Newman were used for all experimental work. Lysogeny broth (LB) agar was used for all plating steps, and agar plates were incubated overnight at 37°C to allow colony growth. MIC assays were conducted in both cation-adjusted Mueller-Hinton broth (caMHB, Sigma Aldrich) and synthetic wound fluid (SWF). SWF comprised 50% vol/vol peptone water (Sigma Aldrich) and 50% vol/vol fetal bovine serum (Gibco), following the recipe of Werthén *et al*. (56). Synthetic wounds were prepared following the method of Werthén *et al*. Briefly, synthetic wound solution was created on ice and comprised 2 mg·ml^−1^ collagen Type 1 (Corning), 0.01% vol/vol acetic acid, 60% vol/vol SWF, and 10 mM sodium hydroxide. 200 μl of synthetic wound solution was added to the wells of a 48-well plate and placed under short-wave UV light for 10 minutes to sterilise each wound. Wounds were then incubated at 37°C for 50-60 minutes to catalyse full polymerisation of the collagen matrix. Wounds were inoculated as follows. Colonies of bacteria from an overnight LB agar plate were added to 5 ml SWF and incubated with shaking for 6 h at 37°C. These starter cultures were diluted to an OD_600_ of 0.05-0.1 and 50 μl of the dilute inoculum was pipetted onto each prepared wound. Biofilms were grown for 24 h at 37°C.

### Vinegar

Commercially-produced vinegars were purchased as detailed in Table 1. The pH of an aliquot of each vinegar was measured using an ETI 8100 pH meter (Electronic Temperature Instruments Ltd). All vinegars were stored in their original bottles in darkness at room temperature.

**Table 1.**
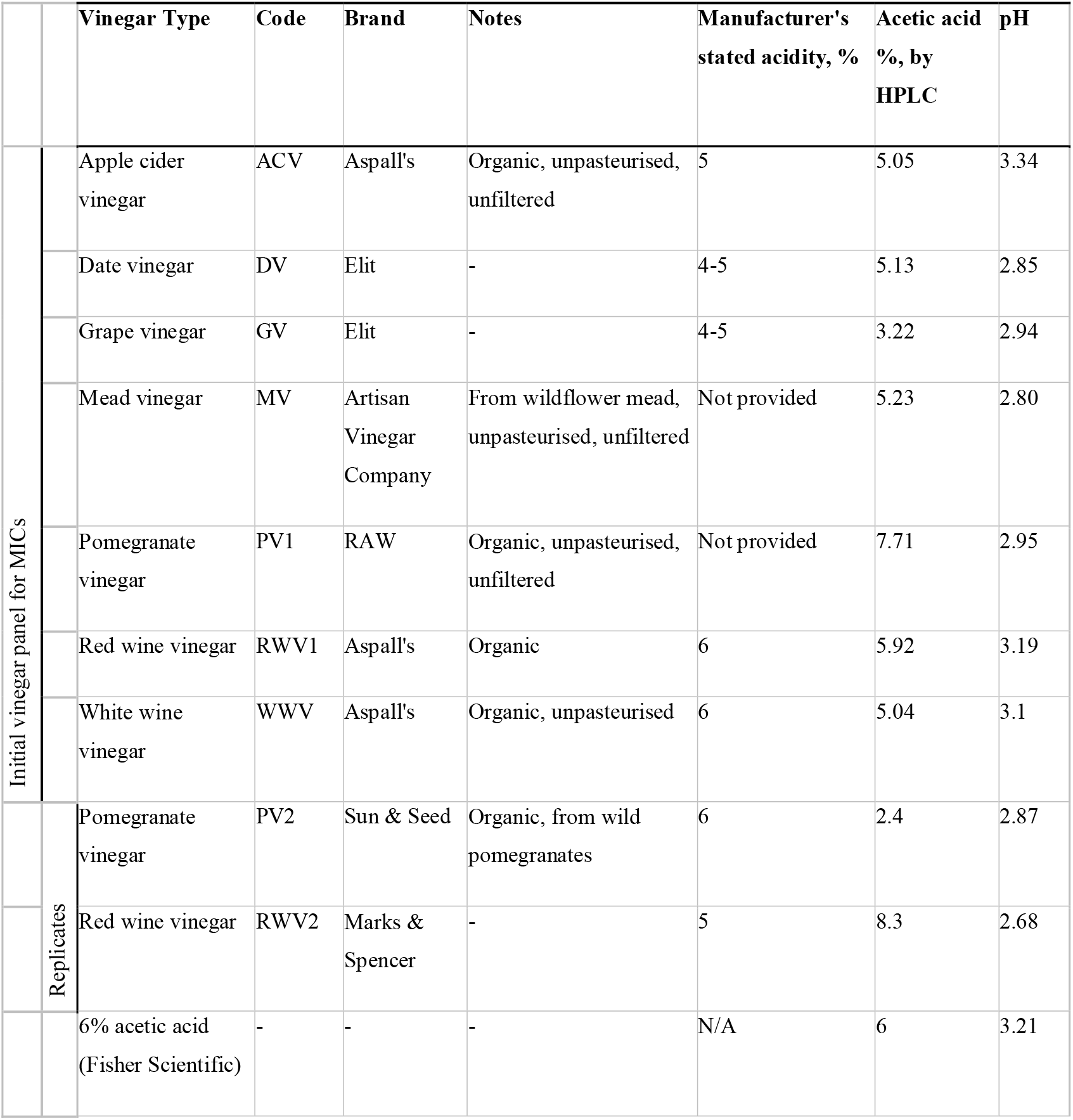
Properties of vinegars and acetic acid.

### Honey

Two commercially-available medical grade honey preparations were used, MediHoney® (product #31805 80% *Leptospermum* honey wound and burn dressing, Dermasciences) and Revamil® (100% honey gel for wound treatment, Oswell Penda). Both of these brands appear on the British National Formulary section on wound management, which lists medicines that may be used by the UK National Health Service (30). Both honeys were stored in their original packaging at room temperature, following manufacturers’ recommendations.

### Analysis of Vinegars by High-Performance Liquid Chromatography (HPLC)

Reversed-phase HPLC was done on an Agilent 1260 infinity II preparative HPLC system; comprising 600 bar quaternary pump, dual-loop aurtosampler, 8-channel UV-based diode array detector (DAD), measuring from 190 to 400nm, and running OpenLab CDS software (version 2.6). Separation was achieved with an Agilent Infinity Lab Poroshell 120 EC-C18 (4.6 × 150mm, 2.5 μm particle size) column First, an undilute aliquot of each vinegar were analysed to capture a chromatographic “fingerprint” of each vinegar, over a 31-minute time period, at 3 wavelengths 210, 260 and 280 nm. A scan was run, and several chromatographic peaks were observed with their peak absorbance near to 260 and/or 280 nm. The method used gradient-grade acetonitrile (ACN; VWR International) and 18.2 MΩ resistivity ultra-pure water (produced from triple-red ultrapure water system) with a starting mix of 5% ACN to 95% water. For 10 minutes a gradient of 5-25% acetonitrile in water was run through the column, after which the concentration gradient was increased from 25-95% and run for a further 21 minutes; both of these steps were run at a flow rate of 0.9 ml·min−1 at ∼240 Bar. An injection volume of 30 μl was used, and separation and detection was done at 25°C. Next, the vinegars were diluted 1/6 in sterile water (Hyclone WFI quality) and triplicate aliquots of each diluted vinegar were run through the HPLC using the same mobile phase and just the first 10 minutes of the above method. The acetic acid concentration was quantified in these aliquots. This dilution factor was chosen because it allowed pure acetic acid at concentrations typical of those found in commercial vinegars to be accurately quantified (i.e. the UV signal was not saturated). The reduction in time to 10 minutes was made because no peaks were observed after 10 minutes in the original 31-minute HPLC run. The injection volume used was 25 μl, and the column temperature was at ∼25°C. This method was also used to analyse a calibration series of 3, 2, 1.5, 1, 0.75, 0.5, 0.38, 0.25, 0.19 and 0.125% vol/vol pure acetic acid (Fisher Scientific) solutions diluted in distilled water (Hyclone WFI quality, Cytiva). A retention time of approximately 2.6 minutes was associated with acetic acid in standard solutions. A calibration curve was created using peak area (mAU^2^) at 210 nm and the calibration equation calculated using OpenLab CDS Data Analysis software (version 2.6).

### Minimum Inhibitory Concentration (MIC) Assays

MIC assays were conducted following CLSI guidelines. 50 μl of each vinegar, or a 6% vol/vol acetic acid solution, was two-fold serially diluted from 100% to 0.098% vol/vol in either caMHB or SWF, in duplicate, in a 96-well microtiter plate (Corning Costar). Five to ten colonies of either *P. aeruginosa* or *S. aureus* were selected from overnight LB agar plates and resuspended in either caMHB or SWF to a 0.5 McFarland standard (OD600 ∼0.08–0.1). The bacterial inoculum was then diluted in the same growth medium to a bacterial density of ∼5×10^5^ colony-forming units (CFU) ·ml^−1^ and 50 μl of this dilution added to all test wells. Antibiotic-free media + bacteria wells, and sterile media wells, were used as growth controls and sterility controls, respectively. The bacterial inoculum was serially diluted in phosphate-buffered saline (PBS,) and plated on LB agar to confirm the bacterial density in the inoculum. 96-well plates were incubated at 37°C for 18 h, and then bacterial growth in each well was visually assessed. The lowest concentration of vinegar or acetic acid with no visible bacterial growth was recorded as the MIC. Minimum bactericidal concentrations (MBCs) were determined by spotting 5 μl from each well of the plate used to determine MICs onto an LB agar plate incubating overnight at 37°C. The lowest treatment concentration with no visible bacterial growth was recorded as the MBC. The duplicate replicates of all MIC or MBC values obtained were within one 2-fold dilution.

### Biofilm Killing Assays in an *In Vitro* Synthetic Wound Model

To assess the dose response of biofilms to selected vinegars or acetic acid, synthetic wound biofilms of either *P. aeruginosa* or *S. aureus* were prepared as above in *Bacterial Strains and Culture Conditions*, and then treated topically with different doses of either water, pure acetic acid, RWV1, RWV2, PV1 or PV2 by pipetting 100 μl of the test substance onto the top of the wound. (See Table 1 for key to vinegar types). Doses were tested in triplicate for each species. RWV1, PV1 and acetic acid, plus water controls, were tested in one experiment. RWV2 and PV2, plus water controls, were tested in a second experiment; the work was divided into two experiments due to the labour-intensive nature of working with the wound model. After 24h exposure to the treatments, the collagen in the wounds was digested to release bacteria by adding 300 μl 0.5 mg·ml^−1^ collagenase type 1 (EMD Millipore Corp, USA) and incubating for 1 h at 37°C. Digested wounds were mixed by pipetting, serially diluted in PBS and plated on LB agar. The number of viable bacteria in each biofilm was estimated using colony counts. To assess the combined effects of selected vinegars or acetic acid + medical-grade honey on biofilms, synthetic wound biofilms of either *P. aeruginosa* or *S. aureus* were prepared as above, and then treated topically with either water; RWV1, PV1, RWV2, PV2 or acetic acid at concentrations corresponding to 0.5% acetic acid; 30% wt/vol MediHoney® 80% gel; 30% wt/vol Revamil® 100% honey gel; or each honey gel combined with each vinegar/acetic acid. In all cases, the total treatment volume was 100 μl, the final concentration of acetic acid was 0.5% and the final concentration of honey gel was 30%. RWV1, PV1 and acetic acid, plus water controls, were tested in one experiment; RWV2 and PV2, plus water controls, were tested in a second experiment. Each treatment was added to triplicate wounds containing *P. aeruginosa* and triplicate wounds containing *S. aureus*. Treated biofilms were incubated at 37°C for 24 h, and then wounds were enzymatically digested, diluted and plated, as in the dose response experiment. The two synergy experiments were replicated using fresh starter cultures of each bacterium.

### Statistical Analyses

For the vinegar or acetic acid dose response experiment, CFU data were recorded and analysed in R 4.0.4 (57) by using the package *drc* (58) to fit a dose response curve for each treatment and calculate the EC_50_. For the synergy experiment, CFU data were recorded and analysed in R 4.0.4. Data were log-transformed to meet the assumptions of general linear modelling and analysed using ANOVA to test for effects of starter culture used, acid treatment, honey treatment and acid treatment*honey treatment. For the dataset testing RWV2 and PV2 and their combinations with honey against *P. aeruginosa* biofilms, one data point was lost and the *car* package (59) was used to conduct the ANOVA using Type II sums of squares to account for non-orthogonality. Planned contrasts using *t*-tests were used to make post-hoc comparisons between the CFU in each treated biofilm and the relevant water-treated control. A significant acid treatment*honey interaction term in the ANOVA was taken as evidence that synergy could exist between at least one acid treatment and one type of medical-grade honey. We used both the Response Additivity and the Bliss Independence models (60) as a null hypotheses for additive interactions between acetic acid or vinegar and medical-grade honey, for reasons explained in the main text. The R package *lsmeans* (61) was used to extract fitted means and 95% confidence intervals for CFU observed in combination treatments, and this was compared with the expected mean CFU under the null hypotheses of Bliss independence as explained in the main text. Graphs were drawn using DataGraph 5.0 (Visual Data Tools, Inc).

## Results

### A systematic review of published evidence for antimicrobial activity of vinegars

A systematic review was performed in November 2021 and repeated in January 2023 to analyse the evidence for differences in antimicrobial properties across vinegar types, and to assess the physiochemical properties that may affect its potency. We searched for papers that included quantitative data on the antimicrobial effect of fermented vinegars against common human bacterial and fungal pathogens and/or against the bacterium *Bacillus subtilis*. As of this date, there were no reviews compiling quantitative evidence.

Full-text articles, as identified following the process outlined in Figure 1 (62), were sorted by quantification method, and papers measuring the minimum inhibitory concentration (MIC, n = 28) or zone of inhibition (ZOI, using either disk- or well-diffusion techniques, n = 49) of vinegars were identified and assessed for key quality parameters. The full list of papers, and summary information, is provided in Table S1. Many papers reporting MIC data of vinegars contained unclear experimental protocols, failing to explicitly state media used, growth conditions and/or method of MIC (i.e., agar dilution or microdilution). Three papers used a non-standard agar dilution method as opposed to microdilution. A number of different media were used during microdilution assays, including Müller-Hinton broth, as endorsed by the CLSI guidelines, tryptic soy and brain heart infusion broths, which are likely to cause some variability in MIC values obtained (63). Reporting of quality parameters, such as number of replicates used, and the presence of appropriate positive and negative controls was poor in the MIC dataset, with only 5 papers in total containing at least two repeats and a negative, untreated, control (i.e. cultures were treated with sterile distilled water).

**Figure 1.**
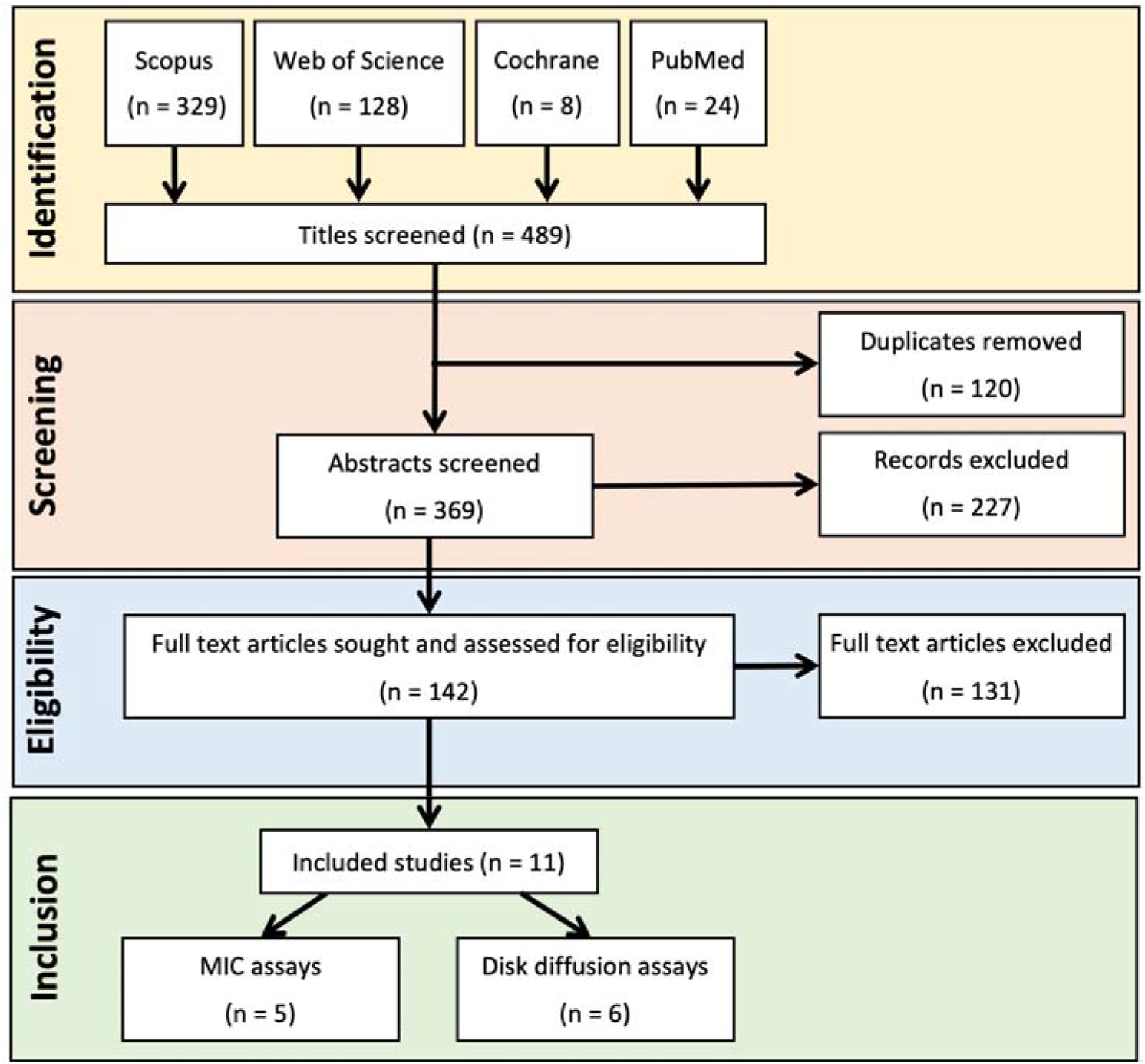
PRISMA diagram showing steps in the systematic review of research articles testing the antimicrobial activity of vinegar. See reference (62) for Preferred Reporting Items for Systematic reviews and Meta-Analyses (PRISMA) guidelines.

Studies using a well-diffusion methodology also lacked sufficient experimental detail. Only three studies employing a well-diffusion technique reported agar depth, a number of these also lacked sufficient replication, and/or details of a negative control. The data is therefore incomparable and was excluded from further analysis. In comparison, studies using the disk diffusion method were overall well standardised, with growth medium, disk size and growth temperature being recorded in the majority of cases. However, the lack of reporting for key details such as number of replicates and the presence of a negative control was still problematic in the disk diffusion dataset. Overall, 6 papers were deemed eligible for full text analysis and were included in the systematic review.

Table S2 contains data extracted from 11 papers included for further analysis with MIC or ZOI data (64-74). These included tests of four Gram-negative bacterial species (*Escherichia coli, Klebsiella pneumoniae, Pseudomonas aeruginosa* and *Salmonella enterica*); four Gram-positive bacterial species (*Bacillus subtilis, Listeria monocytogenes, Staphylococcus aureus, Streptococcus mutans* and *Streptococcus pyogenes*); five named fungal species (*Aspergillus* sp., *Candida albicans, Candida tropicalis, Curvularia* sp., *Lichtheimia* sp.) and an unspecified yeast isolate.

The studies in this review assessed a large range of commercially and traditionally produced vinegars from diverse botanical origins: blueberry, apple, rice, balsamic, red wine, white wine, rose, grape, pomegranate, date, garlic, quince, peach, pineapple, cornelian cherry, purple onion, grape+lavender honey, apple+lavender honey, gilaburu (*Viburnum opulus*) *Eucommia ulmoides* leaves, *Physalis pubescens*, and *Hylocereus monacanthus*. Studies in this review attributed the antimicrobial activity of vinegars to the presence of acetic acid (67, 71, 72), but also suggest that polyphenolic compounds and the presence of other organic acids may play a role in antimicrobial activity (65-67, 71). We cannot conclude from the published literature whether the activity of vinegar is entirely due to its acetic acid content or if whole vinegar is more or less effective than its acetic acid content, i.e. whether other compounds in vinegar potentiate or antagonise the activity of acetic acid. This is because none of the included studies quantified acetic acid in the vinegars used and compared their activity to an equivalent concentration of pure acetic acid. However, the vinegar MICs reported by Kara et al (73) are very low in equivalent % acetic acid, often lower than what is typically reported for pure acetic acid (Table S3). Further, their principal component analysis of vinegar activity and composition showed that MIC does not necessarily correlate with acetic acid content. Due to the variation in methodologies used in the studies included for analysis, it is also challenging to produce meaningful comparisons across studies or to conclude if the antimicrobial activity of vinegar varies significantly by botanical origin. In our previously published assessment of the antimicrobial properties of stinging nettle (*Urtica* spp.) preparations, it was noted that some preparations called for combining nettles with vinegar. Pilot experiments were therefore conducted to assess the MICs and MBCs of several types of commercial vinegars to see if they varied across brands. Red wine vinegar, white wine vinegar and cider vinegar all had low MICs/MBCs for the test species, and these were comparable between species and with MIC/MBC of 6L% acetic acid, although the cider vinegar showed some variability in MIC/MBC between replicates (74). However, a more rigorous laboratory assessment of the activity of different vinegars is clearly needed.

### Characterisation of selected vinegars by high-performance liquid chromatography

We selected seven types of vinegar, commercially produced from a range of starting materials, for investigation. These are summarised in Table 1 and were chosen based on the likely fruits and fruit-based alcoholic beverages used to produce fermented vinegars around the Mediterranean, in Europe and in the Middle East throughout history. Vinegars in our initial panel for MIC testing were analysed by reversed-phase high-performance liquid chromatography (RP-HPLC). We first obtained chromatograms of undiluted vinegar samples to visualise the chromatographic ‘fingerprint’ of the different vinegars, and thenceforth worked with vinegars diluted to bring the expected acetic acid concentration into a range quantifiable using HPLC. Representative chromatograms from triplicate aliquots of each vinegar at 210nm are shown in Figure 2; triplicate chromatograms for each vinegar are provided in Figure S1 and similar chromatograms were obtained at 260nm and 280nm (Figures S2, S3). As can be seen in Figure 2, all vinegars had a clear peak at approx. 2.6 minutes, which was confirmed to be acetic acid by testing against an external acetic acid standard (Figure S4). Other peaks at approx. 1-3 minutes are likely to represent other weak organic acids, while peaks at later times are likely to be polyphenols and other less polar molecules. With the exception of pomegranate vinegar, the samples had very few peaks after approx. 4 minutes, and the non-pomegranate vinegars had qualitatively similar chromatograms. The chromatogram for pomegranate vinegar was much more complex, with multiple, often not clearly resolved, peaks running at times later than 4 minutes. We attempted to use external standards for a selection of expected compounds to identify some of these peaks but, as is often the case with complex mixtures of natural products, we could not unambiguously identify any other peaks in the vinegars. We used an acetic acid calibration series at 210 nm to quantify the acetic acid concentration in the samples. As shown in Table 1, the acetic acid concentration of the vinegars in the initial panel ranged from 3.22% (grape vinegar) to 7.71% (pomegranate vinegar). Figure S4 shows chromatograms from example diluted aliquots of each vinegar used for acetic acid quantification, and a reference standard of acetic acid.

**Figure 2.**
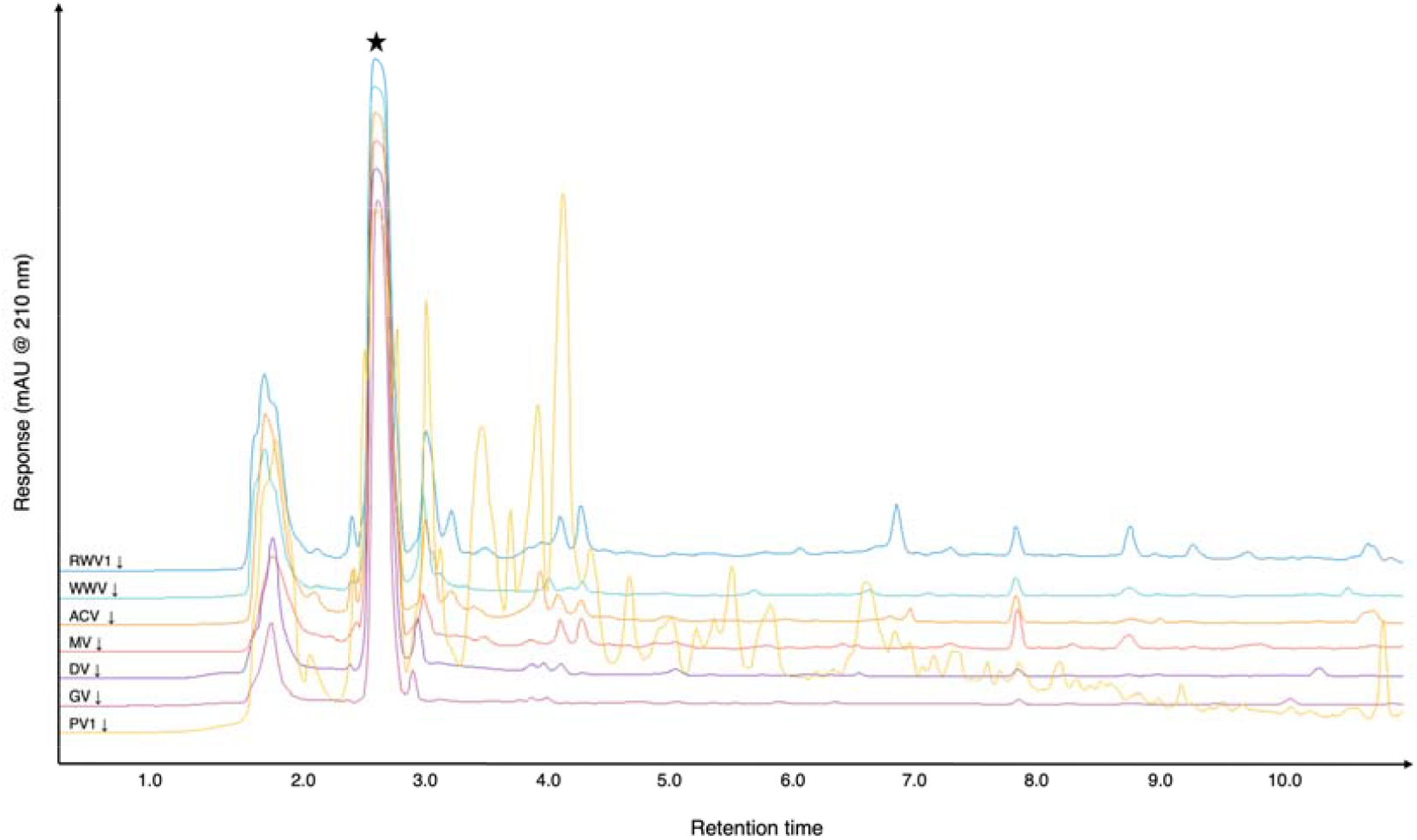
Reversed-phase HPLC chromatograms of diluted vinegar samples at 210nm. Representative chromatograms from triplicate aliquots of selected vinegars; see Table 2 for further details of vinegars tested and Figures S1-S3 for individual chromatograms of aliquots read at 210, 260 and 280 nm. The peak corresponding to acetic acid is marked with a star (see Figure S4).

### An assessment of the antibacterial and antibiofilm activity of different vinegars using *P. aeruginosa* and *S. aureus* as example Gram-negative and Gram-positive bacterial pathogens

We conducted MIC testing by broth microdilution for the initial vinegar panel and pure acetic acid against *P. aeruginosa* PA14 and *S. aureus* Newman in cation-adjusted Muller-Hinton Broth (caMHB) following CLSI guidelines (75). In parallel, we ran the MIC tests in synthetic wound fluid (SWF, (56)), which provides a more *in vivo*-like model of wound exudate than caMHB. This is important because the environment in which bacteria grow can affect both the activity of antimicrobials (e.g. if they bind serum, which is present in wound exudate and in SWF) and the susceptibility of bacteria to antimicrobials (due to physiological changes in gene expression which alter antibiotic targets, metabolism or efflux). MICs can thus be very different in standard media and in an infected host, and using host-mimicking medium can help to close the gap between *in vitro* estimates of susceptibility, and *in vivo* susceptibility (62). We chose SWF as a host-mimicking medium due to the prevalent historical use of vinegar and honey in treatments for wounds and other skin and soft tissue infections. The MICs, initially expressed as a % (vol/vol) of vinegar (Table S4) were converted to an equivalent % acetic acid using the data in Table 1, and the results are presented in Figure 3. Using this adjustment, MICs which are < the MIC of pure acetic acid reflect the vinegar being more antibacterial than expected if acetic acid is the only explanation for their activity, i.e. additional antibacterial molecules and/or molecules which potentiate the activity of acetic acid are present in the vinegar. Conversely, MICs which are > the MIC of pure acetic acid suggest that the vinegar contains molecules which antagonise the activity of acetic acid.

**Table 2.** EC_50_ of pure acetic acid, red wine vinegars and pomegranate vinegars against mature biofilms of *P. aeruginosa* and *S. aureus* grown in a synthetic wound model.

|  | EC<sub>50</sub> (% acetic acid equivalent) |  |
| --- | --- | --- |
|  | *P. aeruginosa* | *S. aureus* |
| Acetic acid (replicate 1) | 0.56 | 0.53 |
| Acetic acid (replicate 2) | 0.49 | 0.33 |
| RWV1 | 0.79 | 0.72 |
| RWV2 | 0.96 | 0.89 |
| PV1 | 0.94 | 0.96 |
| PV2 | 0.28 | 0.01 |

**Figure 3.**
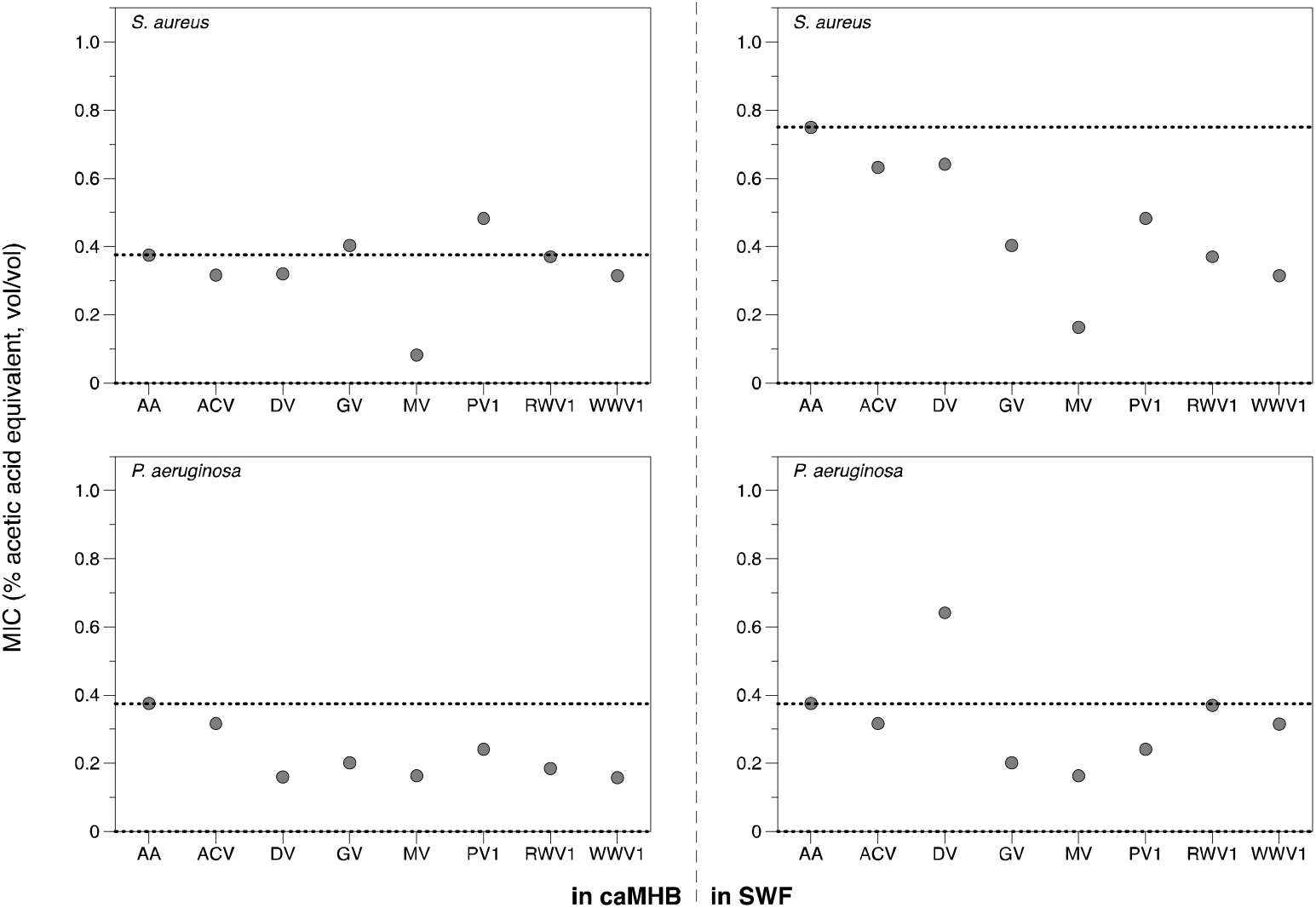
MICs of selected vinegars and acetic acid. MIC tests were conducted against *P. aeruginosa* PA14 and *S. aureus* Newman using a broth microdilution method according to EUCAST guidelines. The assay was repeated in cation-adjusted Muller-Hinton broth (caMHB, left) and synthetic wound fluid (SWF, right).

As shown in Figure 3, in caMHB, most vinegars were about as effective as an equivalent concentration of pure acetic acid against *S. aureus*; the exception was mead vinegar, which was considerably more active than pure acetic acid. However, most of the vinegars were slightly more effective than pure acetic acid against *P. aeruginosa* in caMHB, with the exception of apple cider vinegar, which had very similar activity to that of pure acetic acid. Conducting the MIC assay in SWF had little effect on the activity of vinegars against *P. aeruginosa*, except that date vinegar became less active than expected from its acetic acid content. However, for *S. aureus*, five of the seven vinegars (grape, mead, pomegranate, red wine and white wine vinegars) had a much lower MIC than pure acetic acid when tested in SWF. Taken together, these results show that some vinegars contain antibacterial molecules in addition to acetic acid, and/or molecules that potentiate the activity of acetic acid, depending on the target bacterial species and the test media.

We selected two vinegars for further exploration of their activity against biofilms of *P. aeruginosa* and *S. aureus*. These were red wine vinegar (RWV) and pomegranate vinegar (PV). We chose RWV as a representative of the six vinegars with very similar chromatograms, and because wine was very likely a common starting material for vinegar manufacture in different locations and in different time periods. PV was chosen as its chromatogram was so different from the other vinegars explored, suggesting that it is a more complex preparation. We purchased additional examples of RWV and PV, from different manufacturers, to see if any characteristics of the vinegar types were reproducible when the vinegars were obtained from different sources. In Table 1, the initial RWV and PV used for MIC testing are denoted RWV1 and PV1, and the new samples are denoted RWV2 and PV2. HPLC analysis of undiluted RWV2 and PV2 showed that they had similar chromatograms to RWV1 and PV1, respectively (Figure 4, S5, S6, S7). Like RWV1, RWV2 contained a relatively high concentration of acetic acid (RWV1: 5.92%; RWV2: 8.30%). Interestingly, while PV1 had the second highest concentration of all vinegars tested (7.71%), PV2 had the lowest (2.4%).

**Figure 4.**
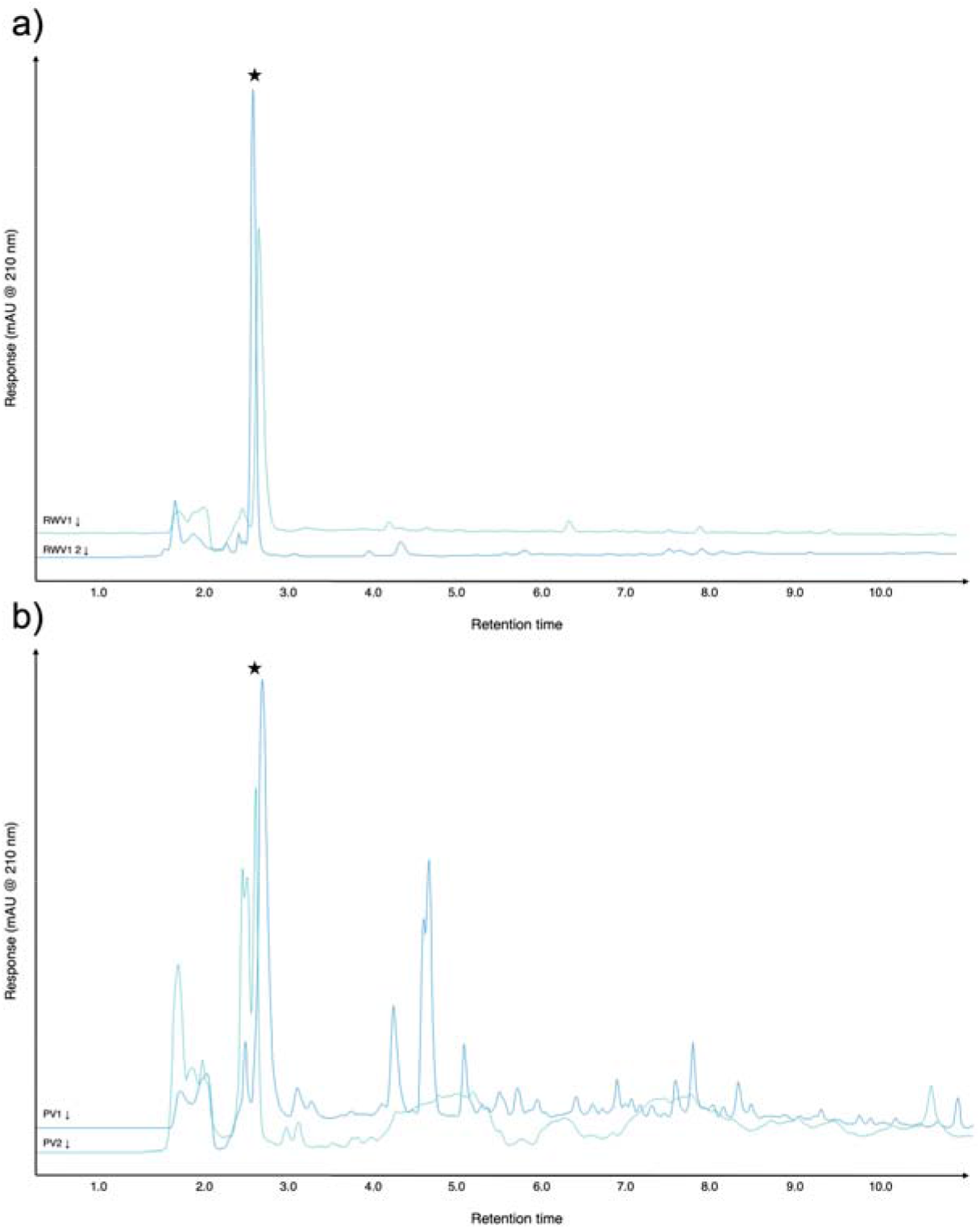
Chromatograms of example aliquots of different red wine and pomegranate vinegars. Representative chromatograms from triplicate aliquots of a) RWV1 and RWV2, b) PV1 and PV2; diluted samples at 210nm. See Figures S5-S7 for individual chromatograms of aliquots read at 210, 260 and 280 nm.

Biofilms of each bacterial species were grown in a SWF-based soft-tissue wound model (55). The model comprises SWF solidified with collagen. Biofilms were allowed to establish in the model for 24 hours, and then treated topically with different doses of water, pure acetic acid, RWV1, RWV2, PV1 or PV2. Doses used were 0.5%, 1% and 3% acetic acid or acetic acid equivalent. After 24h exposure to the treatments, biofilms were enzymatically degraded to release bacteria and the numbers of viable bacteria in each biofilm estimated using colony counts. Dose response curves were fitted to the data and are shown with a linear *y* axis scale in Figure 5.

**Figure 5.**
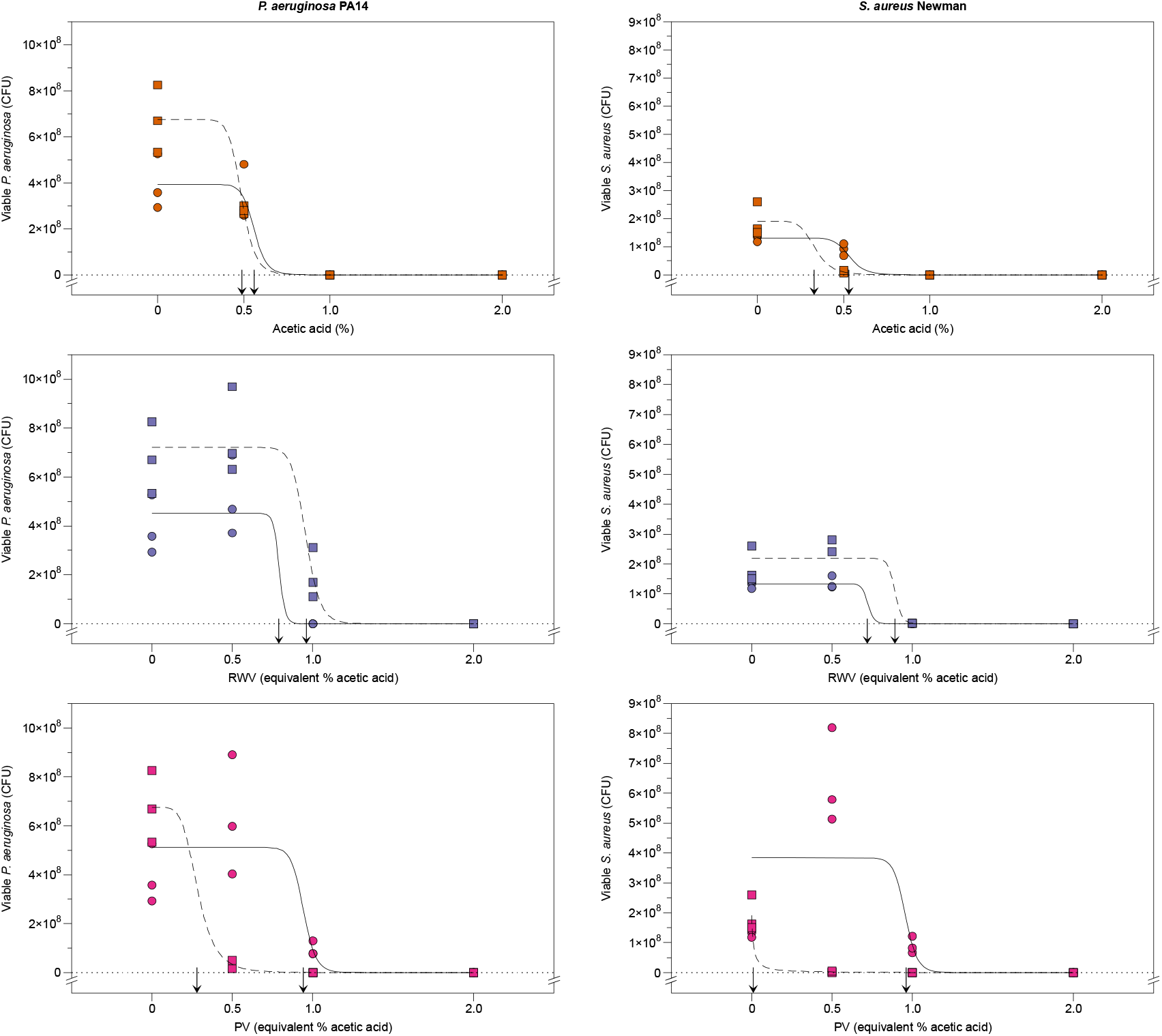
Dose response of mature biofilms to acetic acid, red wine vinegars and pomegranate vinegars. Triplicate synthetic wounds containing mature biofilms of either *P. aeruginosa* PA14 or *S. aureus* Newman were topically treated with water, acetic acid at 0.5%, 1% or 2%, or RWV1, RWV2, PV1 or PV2 at concentrations containing 0.5%, 1% or 2% acetic acid. After 24 hours’ treatment, wounds were enzymatically digested to release bacteria, serially diluted and plated out to count colonies. Colony forming units (CFU) are used to estimate the number of viable bacterial cells in the biofilms. The R package *drc* (58) was used to fit dose response curves for each treatment and the EC_50_ for each treatment is indicated by an arrow at the *x* axis. As The EC_50_ is the concentration predicted to cause a half-maximal kill. Circles and solid lines are used for RWV1 and PV1 data; squares and dashed lines are used for RWV2 and PV2 data. The acetic acid experiment was repeated and data for the two replica experiments are shown with circles+solid lines and squares+dashed lines. These data are provided on a log-transformed *y* axis in Figure S8; raw data and R code are provided in the supplementary data (Document S1).

Raw data are provided plotted on a logged *y* axis in Figure S8. EC_50_ values were calculated for each treatment and are summarised in Table 2. These reveal that RWV1, RWV2 and PV1 are less bactericidal against biofilms of *P. aeruginosa* and *S. aureus* (had higher EC_50_) than equivalent doses of pure acetic acid. This contrasts with the data for MICs in planktonic cultures of the bacteria in SWF, where RWV1 and PV1 were as effective or more effective at inhibiting the bacteria than pure acetic acid. PV2, on the other hand, had a much lower EC_50_ than pure acetic acid against biofilms of both species, reflecting greater biofilm-killing activity.

### Synergistic activity of acetic acid and some vinegars with medical-grade honey, against mature biofilms of *P. aeruginosa* and *S. aureus* in a soft-tissue wound model

We next tested the hypothesis that vinegar and honey may show synergistic antibacterial activity. Our systematic review of the literature did not reveal any published study that tested this hypothesis. We again grew biofilms of either *P. aeruginosa* or *S. aureus* in the synthetic wound model, and aimed to topically treat these with sub-bactericidal concentrations of acetic acid, RWV1, RWV2, PV1 or PV2, alone or in combination with a sub-bactericidal dose of medical-grade honey. Based on the data shown in Figures 5 and S8, we chose 0.5% acetic acid or equivalent concentrations of vinegar. (As PV2 was so active, this dose still caused some killing, but only enough to cause a 2log_10_ drop in colony-forming units; Figure S8). We selected two candidate medical-grade honey ointments: the manuka honey product MediHoney® 80% gel and the non-manuka, peroxide honey product Revamil® 100% gel. Pilot experiments identified a treatment of 30% wt/vol of each honey gel as sub-bactericidal in the biofilm model (data not shown). Acetic acid, RWV1 and PV1 were tested in one set of experiments; RWV2 and PV2 were tested in a second set. Each individual treatment or combination was tested against three biofilms of each species; the experiment was then repeated with a fresh starter culture of bacteria to yield six replicates of each treatment. The results of these experiments are shown in Figure 6.

**Figure 6.**
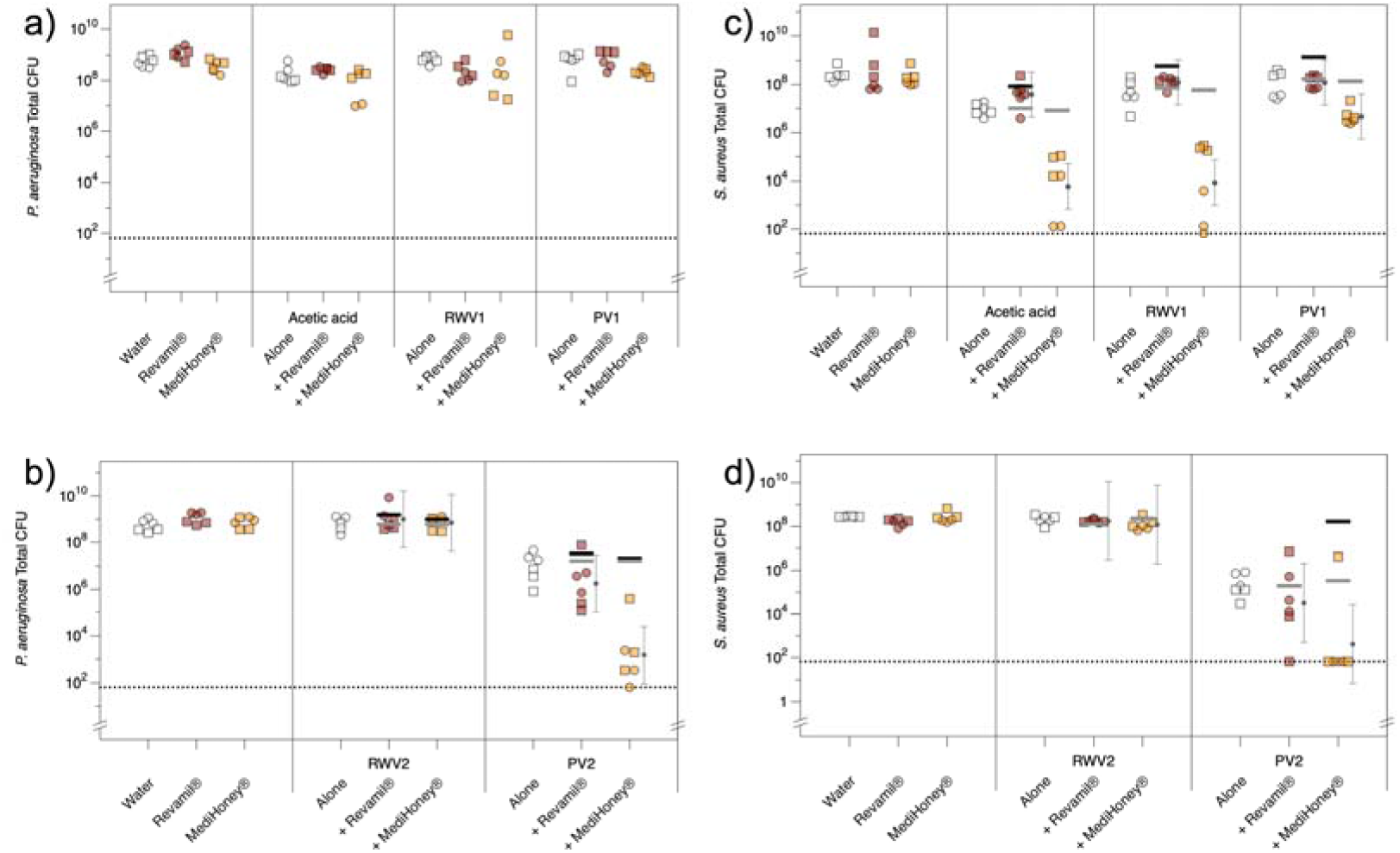
Effect of treating biofilms with acetic acid of vinegar, alone and in combination with medical-grade honey. Triplicate synthetic wounds containing mature biofilms of either *P. aeruginosa* PA14 or *S. aureus* Newman were topically treated with water, 0.5% acetic acid or RWV1, RWV2, PV1 or PV2 at concentrations containing 0.5%, acetic acid, MediHoney® 80% gel at 30% wt/vol, Revamil® 100% gel at 30% wt/vol or combinations of each acid treatment plus each honey treatment. The experiment was repeated with a further set of triplicate wounds from a fresh starter culture. Circles and squares denote the replica experiments. After 24 hours’ treatment, wounds were enzymatically digested to release bacteria, serially diluted and plated out to count colonies. Colony forming units (CFU) are used to estimate the number of viable bacterial cells in the biofilms. **a)** ANOVA revealed no significant interaction between acid treatment and honey treatment (acid F_3,48_=8.35, *p*<0.001; honey F_2,48_=8.18, *p*<0.001; acid*honey F_6,48_=1.46, *p=*0.213). **b)** ANOVA revealed a significant interaction between acid treatment and honey treatment (acid F_2,35_=180, *p*<0.001; honey F_2,35_=21.3, *p*<0.001; acid*honey F_4,35_=20.8 *p*<0.001). Planned contrasts using *t*-tests of each treatment versus the water-treated control showed that only PV2, PV2+Revamil and PV2+Medihoney caused any reduction in CFU compared with the control (*p*<0.001). **c)** ANOVA revealed a significant interaction between acid treatment and honey treatment (acid F_3,48_=83.5, *p*<0.001; honey F_2,48_=179, *p*<0.001; acid*honey F_6,48_=28.2, *p*<0.001). Planned contrasts using *t*-tests of each treatment versus the water-treated control showed that acetic acid, acetic acid + either honey, PV + MediHoney, RWV and RWV+MediHoney caused reductions in CFU compared with the control (*p*≤0.003). **d)** ANOVA revealed a significant interaction between acid treatment and honey treatment (acid F_2,36_=121, *p*<0.001; honey F_2,36_=5.22, *p=*0.010; acid*honey F_4,36_=4.18 *p=*0.007). Planned contrasts using *t*-tests of each treatment versus the water-treated control showed that only PV2, PV2+Revamil and PV2+MediHoney caused reductions in CFU compared with the control (*p<*0.001). For the data shown in panels b-d, the predicted CFU in combination treatments under the assumption of no additive effects of acetic acid/vinegar and honey were calculated as described in the main text and plotted as horizontal bars. The fitted means and associated 95% confidence intervals for the observed CFU in combination treatments were calculated using the R package *lsmeans* REF and added to the plots as small black circles and associated error bars. The predicted mean CFU for combination treatments under Response Additivity and Bliss Independence are shown as thick horizontal black and grey bars, respectively. Where the 95% confidence intervals around observed means do not overlap with these predicted values, the observed CFU is significantly different from that predicted under the respective null model. Raw data and associated R code are provided in the supplementary data (Document S1).; data were log-transformed prior to analysis to meet the assumptions of ANOVA and the R package *car* (59) was used to conduct the ANOVA on the data in panel b because a missing value led to non-orthogonality.

Tests for synergy of antibacterial agents usually rely on a checkerboard design, where multiple concentrations of each agent are tested and fully cross-factored. This allows the MIC or MBC of each agent to be calculated when used alone, and when in combination with the second agent. Using this data, a fractional inhibitory concentration can be calculated and used to determine if the agents show additive, synergistic or antagonistic activity. Alternatively, a synergy landscape can be plotted using a statistical model chosen based on the known mechanisms of action of the agents and regions of concentrations resulting in additive, synergistic or antagonistic activity can be identified (76, 77). The scale of a checkerboard-style experiment when working with five types of acetic acid treatment, two different honeys and a biofilm model is prohibitive, and so we limited ourselves to testing only one concentration of each agent, which was chosen to be sub-bactericidal or minimally bactericidal based on previous experiments. We therefore used the following procedure to test for synergy. First, for each species we used ANOVA to test for the effect on CFU of starter culture, honey treatment (none, Revamil or MediHoney), acid treatment (in experiment 1: none, AA, RWV1 or PV1; in experiment 2: none, RWV2 or PV2) and acid*honey. Synergy or antagonism can only be possible if the interaction term is significant, because a significant interaction means that the effect of the acid treatment depends on the honey treatment, and *vice versa*. In the experiments using acetic acid, RWV1 and PV1, the interaction was not significant for *P. aeruginosa* (Figure 6a) but was significant for *S. aureus* (Figure 6b). In the experiments using RWV2 and PV2, the interaction was significant for both *P. aeruginosa* and *S. aureus* (Figure 6c,d). Thus we concluded that neither honey was synergistic or antagonistic with acetic acid, RWV1 or PV1 against biofilms of *P. aeruginosa*, but that interactions between honey and acetic acid/vinegar should be further explored in the other data sets.

When testing for synergy between antibacterial agents, various null hypotheses that predict the effect of combination treatment in the absence of synergy have been proposed (77). These depend on the mechanisms of action of the two agents being combined. Acetic acid and other weak acids kill bacteria by crossing the cell membrane, collapsing the proton gradient necessary for ATP synthesis and acidifying the cytoplasm, leading to denaturation of proteins and DNA (see (7, 78) and references therein). Honeys similarly have a range of effects on bacteria: all honeys exert some effect via osmotic stress and low pH; peroxide generated by honeys such as Revamil® produce free radicals that damage components of the cell membrane, wall and intracellular targets; and methylglyoxal present in manuka honeys such as MediHoney® can disrupt protein and DNA synthesis, as well as altering cell membrane permeability and damaging cell surface structures (see (79) and references therein). Thus acetic acid (or vinegar) and honey likely attack multiple, partially overlapping sets of cellular targets and do not have identical mechanisms of bactericidal action.

Different reference models of drug synergy make different assumptions about the test agents’ mechanisms of action and dose-response curves. Because our test agents have multiple, partially overlapping mechanisms of action, and because the dose response relationships in our model has not been fully explored, we assessed their activity using two different synergy models that are amenable to use for single-dose experiments: the Response Additivity or Linear Interaction Effect, and Bliss Independence (reviewed by (60)).

Under Response Additivity, the combined effect of non-interacting agents should be the sum of their individual effects: i.e. if A causes an *a*log_10_ kill and B causes a *b*log_10_ kill, Response Additivity should produce an (*a*+*b*)log_10_ kill (60). Synergy would produce a greater than (a+b)log_10_ kill, and antagonism would produce a less than (a+b)log_10_ kill. We therefore calculated the predicted viable CFU count for each combination treatment under Response Additivity, shown as horizontal black bars on the plots in Fig 6. By plotting the 95% confidence interval around the observed mean CFU in each combination treatment, we can see whether the observed CFU is significantly less than or greater than the predicted CFU under Response Additivity. Using this method, we concluded that PV2 showed synergistic activity with MediHoney and Revamil against *P. aeruginosa* biofilms; however, the magnitude of this effect for Revamil was very small and likely not biologically meaningful (Figure 6c). For *S. aureus* biofilms, acetic acid, RWV1, PV1 and PV2 showed synergistic activity with MediHoney, although this effect was very small for PV1. None of the vinegars, nor acetic acid, showed any meaningful synergy with Revamil against *S. aureus* (Fig 6b,d).

Under Bliss independence, if A and B act independently (additively), and the probability of being a cell being killed (fractional change in viable cells) by A is *p(A)* and the probability of being a cell being killed by B is *p(B)*, then independent action results in a fractional decrease in viable cells under combination treatment of *p(A)+p(B)-[p(A)*p(B)]* (60). This leads to the predicted mean CFUs indicated by the horizontal grey bars in Fig 6; under the Bliss null model, there is no major change in our conclusions; the only difference related to the the two cases where Response Additivity showed statistically significant but very slight synergy of (PV1+Revamil in 6c, PV2 + Revamil in 6d); neither of these combinations was synergistic under Bliss independence.

In conclusion, our results show the potential for some vinegars to have antibacterial activity that exceeds that predicted by their acetic acid content alone, but that this depends on the bacterial species being investigated and the growth conditions (media type, planktonic vs. biofilm). Pomegranate vinegars may be a particularly interesting candidate vinegar for further chemical and microbiological study. We also conclude that there is a potential for acetic acid, and some vinegars, to show synergistic antibiofilm effects with manuka honey.

## Discussion

In natural product (NP) research, the protocols for examining chemical composition are often based on looking at a single ingredient, as has been the case with honey and vinegar as individual components, or a single compound purified from a complex natural product. Here our question concerns the potential of synergistic pairing present in a complex ingredient (vinegar) and an ingredient mixture (*oxymel*), in which the summed activity of compounds present may be greater than each individual part (17, 80). The activity of NPs, and NPs in mixtures, are well understood to be impacted by biological variation present in plants as well as the physiological state of the microbes they are used against (reviewed by (81)). One systematic review reported a great degree of variation in values between brands of medical honeys (82). In the case of medical grade manuka honey, calculation of a unique manuka factor (UMF), which is a measure of its antistaphylococcal activity *in vitro* and is well correlated with its total phenolic and methylglyoxal content, is used to overcome product variation (83). However, in the case of whole vinegar, aside from acetic acid, specific contributing antimicrobial molecules and their biocidal activity is largely unknown. Unsurprisingly, previous research has identified that chemical compounds present in vinegar, and the concentrations at which they are found, vary between batches produced (65). Acetic acid, for topical medical applications, has received the most research attention for its evidenced efficacy but also perhaps due to this challenge in assessing whole vinegar and variability in physicochemical properties in such factors as traditionally distilled vinegars versus commercial vinegars, and the botanical origin of the vinegar (84). This added complexity is part of what makes the investigation of NPs more difficult and time-consuming than purified compounds.

Our systematic review found that the published evidence was insufficient to create comparisons of the antimicrobial activities of vinegars, or to draw meaningful conclusions about the contribution of compounds other than acetic acid to the antimicrobial activities of vinegars, due to the variability in methodologies and variation in quality of the studies. Although there is some evidence that acetic acid has a dose-dependent effect on the antimicrobial activity of vinegars, other molecules present, and interactions between the two have not been fully explored. To address gaps highlighted by the review, we characterised selected vinegars by HPLC and assessed their antibacterial and antibiofilm activity using *P. aeruginosa* and *S. aureus* as example Gram-negative and Gram-positive bacterial pathogens.

We found that the concentration of acetic acid in a range of commercial vinegars did not explain their MIC in planktonic culture. Depending on the growth medium used and the target species, many vinegars had a lower MIC than that of an equivalent concentration of pure acetic acid, suggesting that other compounds present in the vinegars could have potentiating or synergistic effects on the activity of acetic acid. Further investigation of the biofilm eradication activity of two red wine vinegars and two pomegranate vinegars in a synthetic wound model concluded that in a biofilm context, the red wine vinegars and one of the pomegranate vinegars were as effective or slightly less effective than equivalent doses of pure acetic acid, but were still able to clear biofilms of both bacterial species at low doses. The second pomegranate vinegar had a stronger biofilm eradication ability than an equivalent concentration of acetic acid. Further work is needed to identify which compounds present in the vinegar are responsible for lending this additional antibiofilm activity.

We then explored the potential for a combination of medical-grade honey and acetic acid or selected vinegars to show synergistic biofilm eradication activity in the synthetic wound model. The scale of a checkerboard-style experiment was prohibitive given the use of a biofilm model, so we instead used single doses of honey or acetic acid / vinegar that caused no or low levels of killing in the biofilm model. Using the assumption of Response Additivity as a null model, we found that strong synergy existed between manuka-based MediHoney® and three test treatments (acetic acid, one red wine vinegar and one pomegranate vinegar) when used to treat *S. aureus* biofilms. When used to treat *P. aeruginosa* biofilms, only one pomegranate vinegar and the MediHoney® showed strong synergy. None of the vinegars, nor acetic acid, showed any synergy with a peroxide-generating honey preparation under the Response Additivity model for estimating synergy in single-dose experiments. Using Bliss Independence as a null model for synergy testing did, however, suggest some synergy between the peroxide-generating Revamil honey and acetic acid or vinegar.

Further work is needed to fully explore the interactions between acetic acid or vinegars, with manuka honey. Ideally, a range of lab and clinical isolates of *S. aureus* and *P. aeruginosa* should be used in checkerboard and time-kill analysis of honey and acetic acid / vinegar combinations to provide ‘gold standard’ analysis of their interactions and the potential for synergy at a range of doses (76), compared with investigations into the mechanisms by which bacteria treated with combined *versus* individual agents are killed.

Our work confirms earlier suggestions that compounds present in some vinegars may enhance their antibacterial activity above that provided by acetic acid. To the best of our knowledge, our work also provides the first published study to address whether combinations of acetic acid or vinegar with honey can have synergistic antibiofilm effects. We demonstrated that acetic acid can synergise with a manuka-based medical grade honey preparation in a biofilm eradication assay using either *S. aureus* or *P. aeruginosa*. We also found that one pomegranate vinegar showed synergy with the honey against both study bacteria, and that one red wine vinegar showed synergy with the honey against *S. aureus* only.

The results from this preliminary study suggest that future research should concentrate on expanding the investigation of pomegranate vinegar to assess the spectrum of activity against other pathogenic species not included in the present study. This work would support established applications and other ongoing investigations of honey and vinegar / acetic acid in a clinical context. A combined wound dressing product, including both manuka honey and acetic acid, could be a suitable candidate for progression to a Phase I trial to assess superiority to dressings using each agent alone.

## Author Contributions

Conceptualisation: EC

Formal Analysis: FH

Funding Acquisition: EC,FH

Investigation: AB, CdW, FH

Supervision: EC, FH

Writing – original draft: EC, FH

Writing – review & editing: AB, CdW, EC, FH

## Acknowledgements

We thank Naseeba Alozaibi, Arwa Hasan, and Derar Hasan Abdel-Qader for providing translations of the paper by Dallee Hamad into English; Hannah Bower for sharing her manuscript images; Bridget Penman and Oluwatosin Orababa for helpful discussions of using null models to test for synergy in single-dose experiments; the WISB Research Technology Facility, School of Life Sciences, University of Warwick for training and support in the use of Agilent 1260 infinity II Preparative HPLC system; and the Media Preparation Facility in the School of Life Sciences at the University of Warwick, with special thanks to Cerith Harries and Caroline Stewart.

## References

1. Kuropatnicki AK, Klósek M, Kucharzewski M. Honey as medicine: historical perspectives. Journal of Apicultural Research. 2018;57(1):113–8. 10.1080/00218839.2017.1411182

2. Eteraf-Oskouei T, Najafi M. Traditional and modern uses of natural honey in human diseases: a review. Iran J Basic Med Sci. 2013;16(6):731–42.

3. Mazza S, Murooka Y. Vinegars Through the Ages. In: Solieri L, Giudici P, editors. Vinegars of the World. Milano: Springer Milan; 2009. p. 17–39.

4. Krug I. The Wounded Soldier: Honey and Late Medieval Military Medicine. In: Krug I, Tracy L, DeVries K, editors. Wounds and Wound Repair in Medieval Culture: Brill; 2015. p. 194–214.

5. Zargaran A, Zarshenas MM, Mehdizadeh A, Mohagheghzadeh A. Oxymel in medieval Persia. Pharm Hist (Lond). 2012;42(1):11–3.

6. Lylye of Medicynes, Oxford Bodleian Library, MS. Ashmole 1505, fols 147v, 157r.

7. Halstead FD, Rauf M, Moiemen NS, Bamford A, Wearn CM, Fraise AP, et al. The Antibacterial activity of acetic acid against biofilm-producing pathogens of relevance to burns patients. PLOS ONE. 2015;10(9):e0136190. 10.1371/journal.pone.0136190

8. Halstead FD, Rauf M, Bamford A, Wearn CM, Bishop JRB, Burt R, et al. Antimicrobial dressings: Comparison of the ability of a panel of dressings to prevent biofilm formation by key burn wound pathogens. Burns. 2015;41(8):1683–94. https://doi.org/10.1016/j.burns.2015.06.005

9. Bjarnsholt T, Alhede M, Jensen PØ, Nielsen AK, Johansen HK, Homøe P, et al. Antibiofilm properties of acetic acid. Advances in Wound Care. 2015;4(7):363–72. 10.1089/wound.2014.0554

10. Ryssel H, Kloeters O, Germann G, Schäfer T, Wiedemann G, Oehlbauer M. The antimicrobial effect of acetic acid—An alternative to common local antiseptics? Burns. 2009;35(5):695–700. https://doi.org/10.1016/j.burns.2008.11.009

11. Ryssel H, Germann G, Riedel K, Reichenberger M, Hellmich S, Kloeters O. Suprathel–acetic acid matrix versus Acticoat and Aquacel as an antiseptic dressing: an in vitro study. Annals of Plastic Surgery. 2010;65(4).

12. Chen H, Chen T, Giudici P, Chen F. Vinegar functions on health: constituents, sources, and formation mechanisms. Comprehensive Reviews in Food Science and Food Safety. 2016;15(6):1124–38. https://doi.org/10.1111/1541-4337.12228

13. Boban N, Tonkic M, Budimir D, Modun D, Sutlovic D, Punda-Polic V, et al. Antimicrobial effects of wine: separating the role of polyphenols, ph, ethanol, and other wine components. Journal of Food Science. 2010;75(5):M322–M6. 10.1111/j.1750-3841.2010.01622.x

14. ISRCTN11636684: A clinical trial looking at the efficacy and optimal dose of acetic acid in burn wound infections. https://doi.org/10.1186/ISRCTN11636684

15. NIHR SRMRC, AceticA. Examining the efficacy and optimal dose of acetic acid to treat colonised burns wounds IRAS ID 142301; https://srmrc.nihr.ac.uk/trials/acetica/.

16. Nour S, Reid G, Sathanantham K, Mackie I. Acetic acid dressings used to treat pseudomonas colonised burn wounds: A UK national survey. Burns. 2022;48(6):1364–7. https://doi.org/10.1016/j.burns.2021.07.011

17. Caesar LK, Cech NB. Synergy and antagonism in natural product extracts: when 1 + 1 does not equal 2. Natural Product Reports. 2019;36(6):869–88. 10.1039/C9NP00011A

18. Carneiro A, Couto JA, Mena C, Queiroz J, Hogg T. Activity of wine against Campylobacter jejuni. Food Control. 2008;19(8):800–5. https://doi.org/10.1016/j.foodcont.2007.08.006

19. Møretrø T, Daeschel MA. Wine is bactericidal to foodborne pathogens. Journal of Food Science. 2004;69(9):M251–M7. https://doi.org/10.1111/j.1365-2621.2004.tb09938.x

20. Budak NH, Aykin E, Seydim AC, Greene AK, Guzel-Seydim ZB. Functional properties of vinegar. Journal of Food Science. 2014;79(5):R757–R64. https://doi.org/10.1111/1750-3841.12434

21. Kelebek H, Kadiroğlu P, Demircan NB, Selli S. Screening of bioactive components in grape and apple vinegars: Antioxidant and antimicrobial potential. Journal of the Institute of Brewing. 2017;123(3):407–16. https://doi.org/10.1002/jib.432

22. Maddocks SE, Jenkins RE. Honey: a sweet solution to the growing problem of antimicrobial resistance? Future Microbiology. 2013;8(11):1419–29. 10.2217/fmb.13.105

23. Roberts A, Brown H, Jenkins R. On the antibacterial effects of manuka honey: mechanistic insights. Research and Reports in Biology. 2015;6:215–44.

24. Molan PC. Potential of honey in the treatment of wounds and burns. American Journal of Clinical Dermatology. 2001;2(1):13–9. 10.2165/00128071-200102010-00003

25. Food and Drug Administration Executive Summary: Classification of Wound Dressings Combined with Drugs, Prepared for the Meeting of the General and Plastic Surgery Devices Advisory Panel September 20-21, 2016, https://www.fda.gov/media/100005/download.

26. Derwin R, Patton D, Avsar P, Strapp H, Moore Z. The impact of topical agents and dressing on pH and temperature on wound healing: A systematic, narrative review. International Wound Journal. 2022;19(6):1397–408. https://doi.org/10.1111/iwj.13733

27. Jull AB, Cullum N, Dumville JC, Westby MJ, Deshpande S, Walker N. Honey as a topical treatment for wounds. Cochrane Database of Systematic Reviews. 2015(3). 10.1002/14651858.CD005083.pub4

28. Grego E, Robino P, Tramuta C, Giusto G, Boi M, Colombo R, et al. Evaluation of antimicrobial activity of Italian honey for wound healing application in veterinary medicine. Schweizer Archiv fur Tierheilkunde. 2016;158(7):521–7. 10.17236/sat00075

29. Fidaleo M, Zuorro A, Lavecchia R. Antimicrobial activity of some Italian honeys against pathogenic bacteria. Chemical Engineering Transactions 2011. p. 1015–20.

30. British National Formulary https://bnf.nice.org.uk/wound-management/honey-dressings.html [

31. Oxford Health NHS Foundation Trust. Medical honey simplified. A guide to the role of honey in wound management for healthcare professionals, OP-061. 2015.

32. Mitchell T. Use of Manuka honey for autolytic debridement in necrotic and sloughy wounds. J Community Nurs. 2018;32(4):38–43.

33. Cambridge University Library, MS. kk.6.33, fol. 31v.

34. Oxford Bodleian Library, MS. Ashmole 1505, fol. 26r.

35. Oxford Bodleian Library, MS. Ashmole 1505, fol. 27r.

36. Oxford Bodleian Library, MS. Add. A. 106, fol. 183v.

37. Oxford Bodleian Library, MS. Ashmole 1505, fol. 28r.

38. Oxford Bodleian Library, MS. Ashmole 1505, fol. 137v.

39. Cambridge University Library, MS. kk.6.33, fol. 49r.

40. Oxford Bodleian Library, MS. Add. A. 106, fol. 186r.

41. Oxford Bodleian Library, MS. Add. B. 60, fols 101r–101v.

42. Oxford Bodleian Library, MS. Ashmole 1432, fol. 43r.

43. Cambridge University Library, MS. kk.6.33, fols 7r, 80v, 101r.

44. British Library, MS. Lansdowne 680, fol. 62r.

45. Oxford Bodleian Library, MS. Add. A. 106, fol. 285r.

46. Oxford Bodleian Library, MS. Add. B. 60, fol. 29r.

47. British Library, MS. Lansdowne 680, fol. 3r.

48. von Fleischhacker R. Lanfrank’s Science of Cirurgie. London: Pub. for the Early English society b K. Paul, Trench, Trübner & Co. (p. 228); 1894.

49. Oxford Bodleian Library, MS. Douce 304, fol. 19v

50. Connelly E, del Genio CI, Harrison F. Data mining a medieval medical text reveals patterns in ingredient choice that reflect biological activity against infectious agents. mBio. 2020;11(1):e03136–19. doi:10.1128/mBio.03136-19

51. Dallee Hamad B. Inhibitory effect of vinegar and its mixture with honey on some pathogenic bacteria. Journal of Education and Science. 2010;23(4):31–41. 10.33899/edusj.2010.58454

52. nimrouzi M, Abolghasemi J, Sharifi MH, Nasiri K, Akbari A. Thyme oxymel by improving of inflammation, oxidative stress, dyslipidemia and homeostasis of some trace elements ameliorates obesity induced by high-fructose/fat diet in male rat. Biomedicine & Pharmacotherapy. 2020;126:110079. https://doi.org/10.1016/j.biopha.2020.110079

53. Nejatbakhsh F, Karegar-Borzi H, Amin G, Eslaminejad A, Hosseini M, Bozorgi M, et al. Squill Oxymel, a traditional formulation from Drimia Maritima (L.) Stearn, as an add-on treatment in patients with moderate to severe persistent asthma: A pilot, triple-blind, randomized clinical trial. Journal of Ethnopharmacology. 2017;196:186-92. https://doi.org/10.1016/j.jep.2016.12.032

54. IRCT201705261165N20, Effect of squill oxymel in control of mild to moderate fatty liver, http://en.irct.ir/trial/319.

55. Mohammadi-Araghi M, Eslaminejad A, Karegar-Borzi H, Mazloomzadeh S, Nejatbakhsh F. An add-on treatment for moderate COPD with squill-oxymel (a traditional formulation from Drimia maritima (L.) Stearn): A pilot randomized triple-blinded placebo-controlled clinical trial. Evid Based Complement Alternat Med. 2022;2022:5024792. 10.1155/2022/5024792

56. Werthén M, Henriksson L, Jensen PØ, Sternberg C, Givskov M, Bjarnsholt T. An in vitro model of bacterial infections in wounds and other soft tissues. APMIS. 2010;118(2):156–64. 10.1111/j.1600-0463.2009.02580.x

57. R Core Team. R: A Language and Environment for Statistical Computing. R Foundation for Statistical Computing, Vienna, Austria. http://www.R-project.org. 2021.

58. Ritz C, Baty F, Streibig JC, Gerhard D. Dose-response analysis using R. PLOS ONE. 2016;10(12):e0146021. 10.1371/journal.pone.0146021

59. Fox J, Weisberg S. An R Companion to Applied Regression. 2011.

60. Duarte D, Vale N. Evaluation of synergism in drug combinations and reference models for future orientations in oncology. Current Research in Pharmacology and Drug Discovery. 2022;3:100110. https://doi.org/10.1016/j.crphar.2022.100110

61. Lenth RV. Least-squares means: The R package lsmeans. Journal of Statistical Software. 2016;69(1):1 33. 10.18637/jss.v069.i01

62. Page MJ, McKenzie JE, Bossuyt PM, Boutron I, Hoffmann TC, Mulrow CD, et al. Updating guidance for reporting systematic reviews: development of the PRISMA 2020 statement. Journal of Clinical Epidemiology. 2021. https://doi.org/10.1016/j.jclinepi.2021.02.003

63. Ersoy SC, Heithoff DM, Barnes LV, Tripp GK, House JK, Marth JD, et al. Correcting a fundamental flaw in the paradigm for antimicrobial susceptibility testing. EBioMedicine. 2017;20:173–81. 10.1016/j.ebiom.2017.05.026

64. Rabeea IS, Janabi AM. Antibacterial Activity of Different Concentrations of Date Vinegar in Comparison to Ciprofloxacin against Multidrug-Resistance Pseudomonas aeruginosa isolated from infected burn. Anti-Infective Agents. 2018;16:96–9.

65. Kara M, Assouguem A, Kamaly OMA, Benmessaoud S, Imtara H, Mechchate H, et al. The impact of apple variety and the production methods on the antibacterial activity of vinegar samples. Molecules (Basel, Switzerland). 2021;26(18). 10.3390/molecules26185437

66. Kadiroglu P. FTIR spectroscopy for prediction of quality parameters and antimicrobial activity of commercial vinegars with chemometrics. Journal of the Science of Food and Agriculture. 2018;98(11):4121–7. https://doi.org/10.1002/jsfa.8929

67. Jia CF, Yu WN, Zhang BL. Manufacture and antibacterial characteristics of Eucommia ulmoides leaves vinegar. Food Sci Biotechnol. 2020;29(5):657–65. 10.1007/s10068-019-00712-7

68. Darus F, Misa N, Ismail Z, Mahidin H. Assessment of antifungal agent for the treatment of Culvularia sp. and Lichtheimia sp. IOP Conference Series: Earth and Environmental Science. 2019;373. 10.1088/1755-1315/373/1/012019

69. Fonseca M, Santos V, Calegari G, Dekker R, Barbosa-Dekker A, Cunha M. Blueberry and honey vinegar: successive batch production, antioxidant potential and antimicrobial ability. Brazilian Journal of Food Technology. 2018;21:e2017101. 10.1590/1981-6723.10117

70. Pedroso JdF, Sangalli J, Brighenti FL, Tanaka MH, Koga-Ito CY. Control of bacterial biofilms formed on pacifiers by antimicrobial solutions in spray. International Journal of Paediatric Dentistry. 2018;28(6):578–86. https://doi.org/10.1111/ipd.12413

71. Fernandes ACF, de Souza AC, Ramos CL, Pereira AA, Schwan RF, Dias DR. Sensorial, antioxidant and antimicrobial evaluation of vinegars from surpluses of physalis (Physalis pubescens L.) and red pitahaya (Hylocereus monacanthus). Journal of the Science of Food and Agriculture. 2019;99(5):2267–74. https://doi.org/10.1002/jsfa.9422

72. Antoniewicz J, Jakubczyk K, Kwiatkowski P, Maciejewska-Markiewicz D, Kochman J, Rebacz-Maron E, et al. Analysis of antioxidant capacity and antimicrobial properties of selected polish grape vinegars obtained by spontaneous fermentation. Molecules (Basel, Switzerland). 2021;26(16). 10.3390/molecules26164727

73. Kara M, Assouguem A, Fadili ME, Benmessaoud S, Alshawwa SZ, Kamaly OA, et al. Contribution to the evaluation of physicochemical properties, total phenolic content, antioxidant potential, and antimicrobial activity of vinegar commercialized in Morocco. Molecules (Basel, Switzerland). 2022;27(3):770.

74. Harrison F, Furner-Pardoe J, Connelly E. An assessment of the evidence for antibacterial activity of stinging nettle (Urtica dioica) extracts. Access Microbiology. 2022;4(3). https://doi.org/10.1099/acmi.0.000336

75. Clinical Laboratory Standards Institute. Performance standards for antimicrobial susceptibility testing (CLSI document M100, 2020). 2020.

76. Doern CD. When does 2 plus 2 equal 5? A review of antimicrobial synergy testing. J Clin Microbiol. 2014;52(12):4124–8. 10.1128/jcm.01121-14

77. Ianevski A, He L, Aittokallio T, Tang J. SynergyFinder: a web application for analyzing drug combination dose–response matrix data. Bioinformatics. 2017;33(15):2413–5. 10.1093/bioinformatics/btx162

78. Kundukad B, Udayakumar G, Grela E, Kaur D, Rice SA, Kjelleberg S, et al. Weak acids as an alternative anti-microbial therapy. Biofilm. 2020;2:100019. https://doi.org/10.1016/j.bioflm.2020.100019

79. Bouzo D, Cokcetin NN, Li L, Ballerin G, Bottomley AL, Lazenby J, et al. Characterizing the mechanism of action of an ancient antimicrobial, manuka honey, against Pseudomonas aeruginosa using modern transcriptomics. mSystems. 2020;5(3):e00106–20. doi:10.1128/mSystems.00106-20

80. Tyers M, Wright GD. Drug combinations: a strategy to extend the life of antibiotics in the 21st century. Nature Reviews Microbiology. 2019;17(3):141–55. 10.1038/s41579-018-0141-x

81. Connelly E, Lee C, Furner-Pardoe J, del Genio CI, Harrison F. A case study of the Ancientbiotics collaboration. Patterns. 2022;3(12). 10.1016/j.patter.2022.100632

82. Nolan VC, Harrison J, Wright JEE, Cox JAG. Clinical significance of manuka and medical-grade honey for antibiotic-resistant infections: a systematic review. Antibiotics. 2020;9(11):766.

83. Johnston M, McBride M, Dahiya D, Owusu-Apenten R, Nigam PS. Antibacterial activity of Manuka honey and its components: An overview. AIMS Microbiol. 2018;4(4):655–64. 10.3934/microbiol.2018.4.655

84. Arvaniti OS, Mitsonis P, Siorokos I, Dermishaj E, Samaras Y. The physicochemical properties and antioxidant capacities of commercial and homemade Greek vinegars. Acta Sci Pol Technol Aliment. 2019;18(3):225–34. 10.17306/j.Afs.0669

