## Supplementary Figures for "Sweet and sour synergy: exploring the antibacterial and antibiofilm activity of acetic acid and vinegar combined with medical-grade honeys"

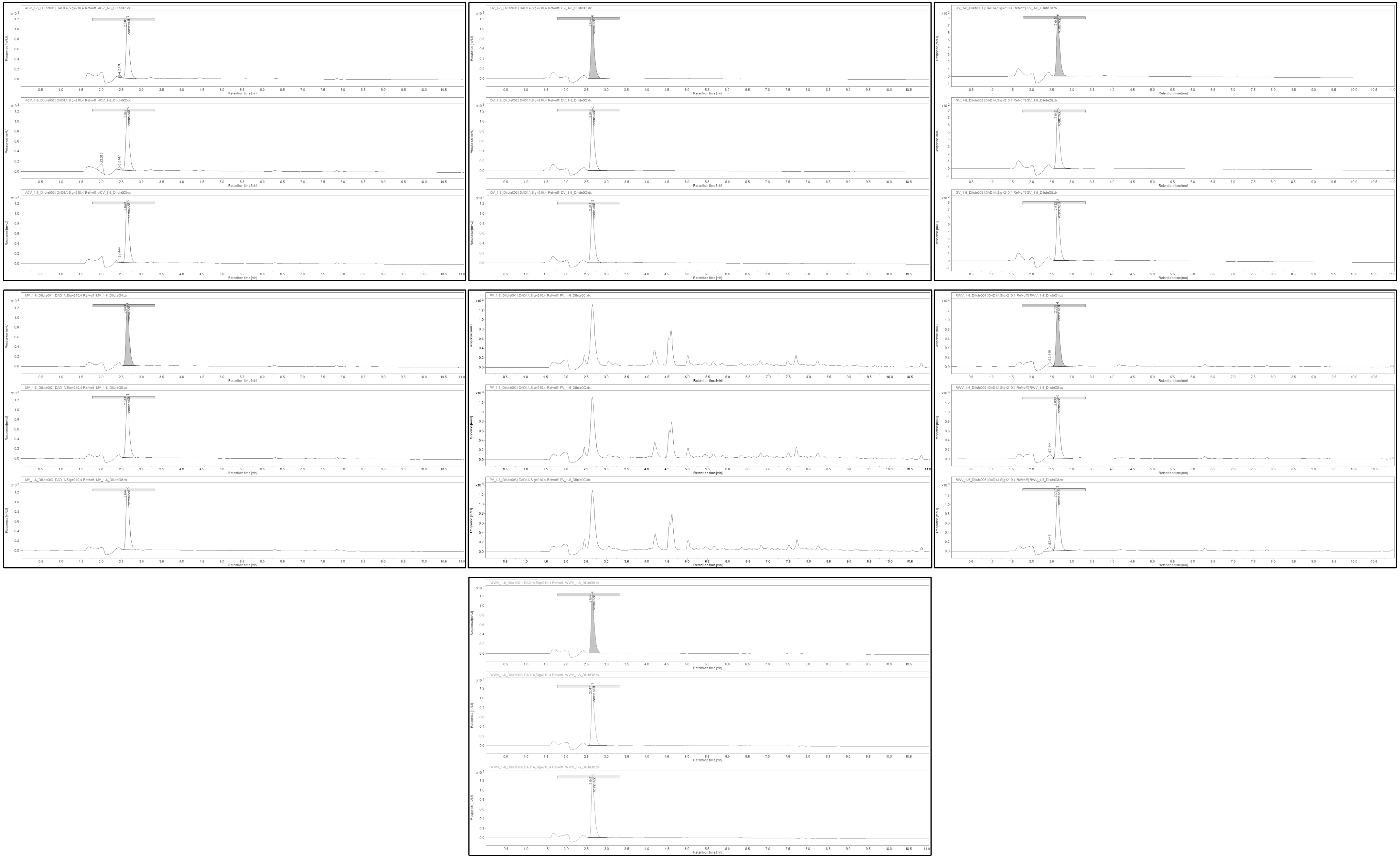

**Figure S1.** Reversed-phase HPLC chromatograms of triplicate, diluted samples of each vinegar from the initial panel, at 210nm

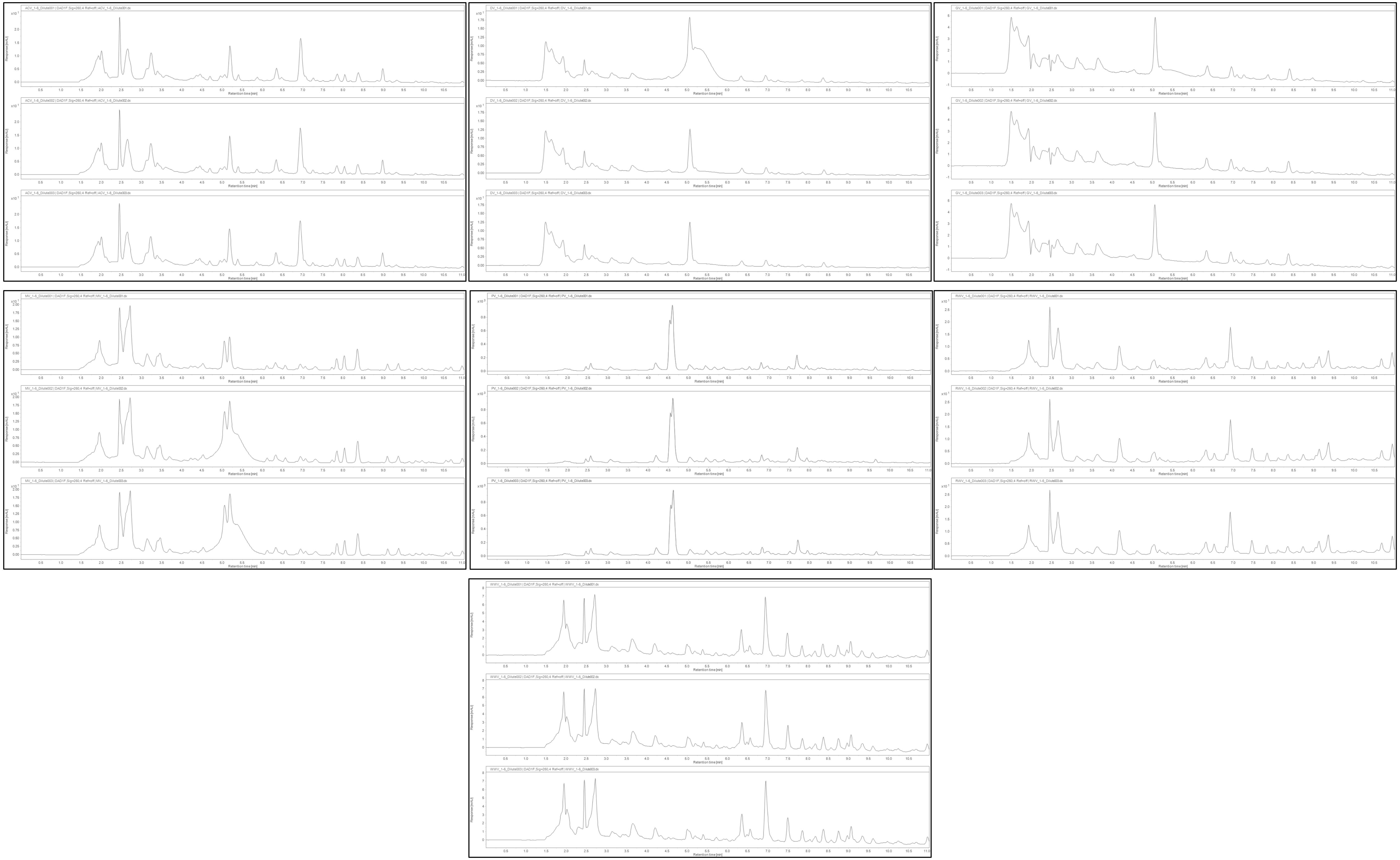

**Figure S1.** Reversed-phase HPLC chromatograms of triplicate, diluted samples of each vinegar from the initial panel, at 260nm

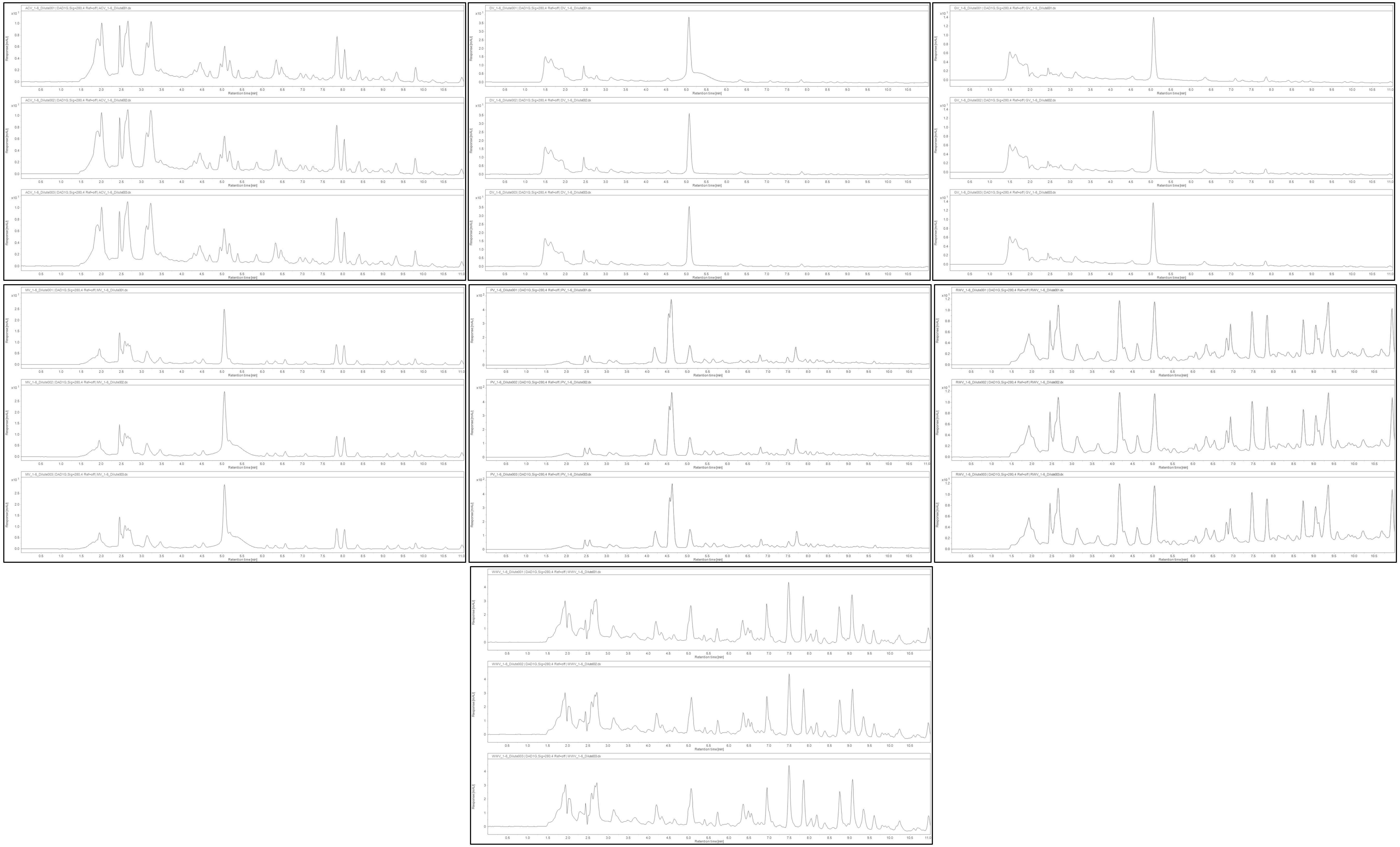

**Figure S3.** Reversed-phase HPLC chromatograms of triplicate, diluted samples of each vinegar from the initial panel, at 280nm

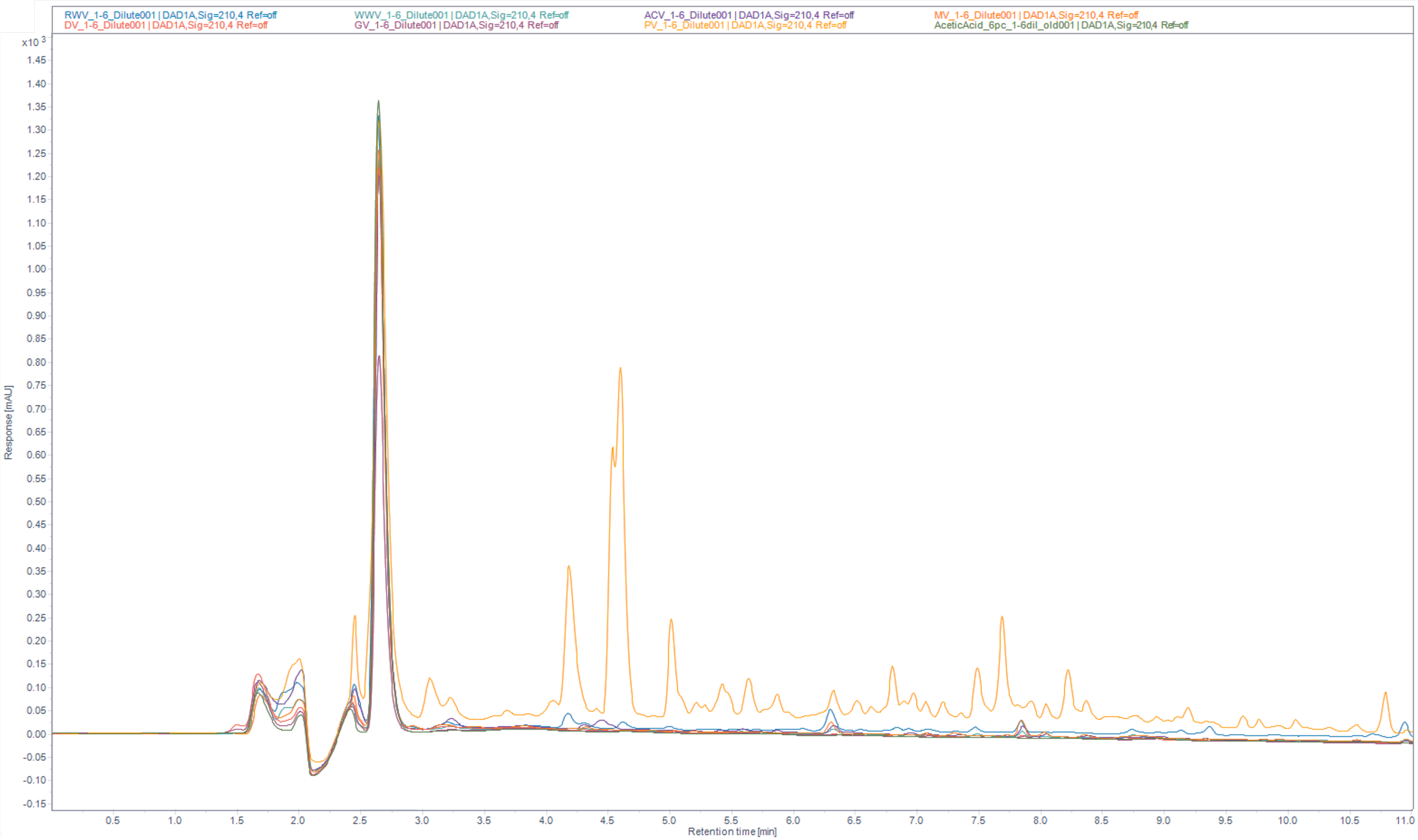


**Figure S4.** Reversed-phase HPLC chromatograms of diluted samples of each vinegar from the initial panel, plus an acetic acid standard, at 210nm.

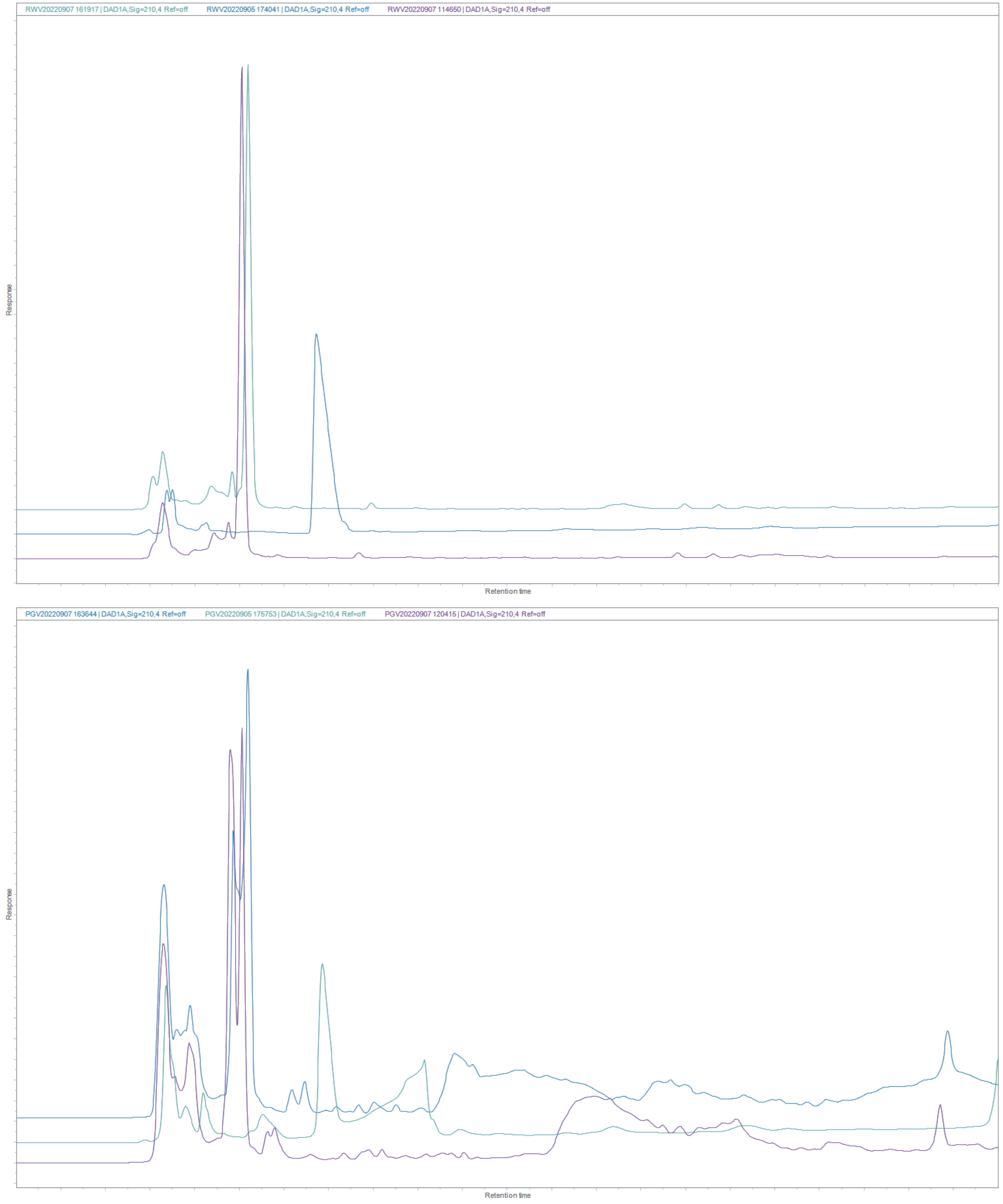


**Figure S5.** Reversed-phase HPLC chromatograms of triplicate samples of RWV2 and PV2, at 210nm

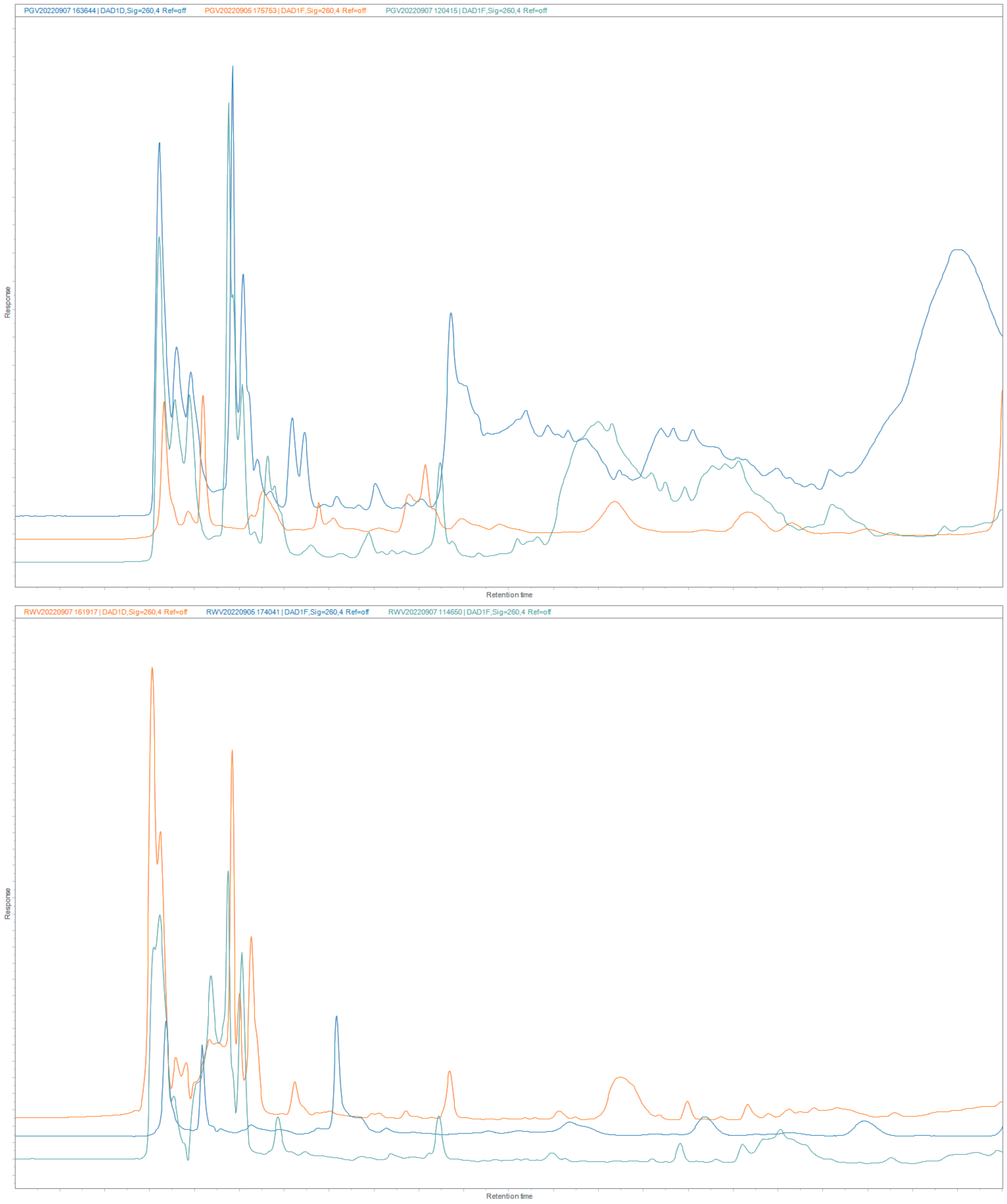


**Figure S6.** Reversed-phase HPLC chromatograms of triplicate samples of RWV2 and PV2, at 260nm

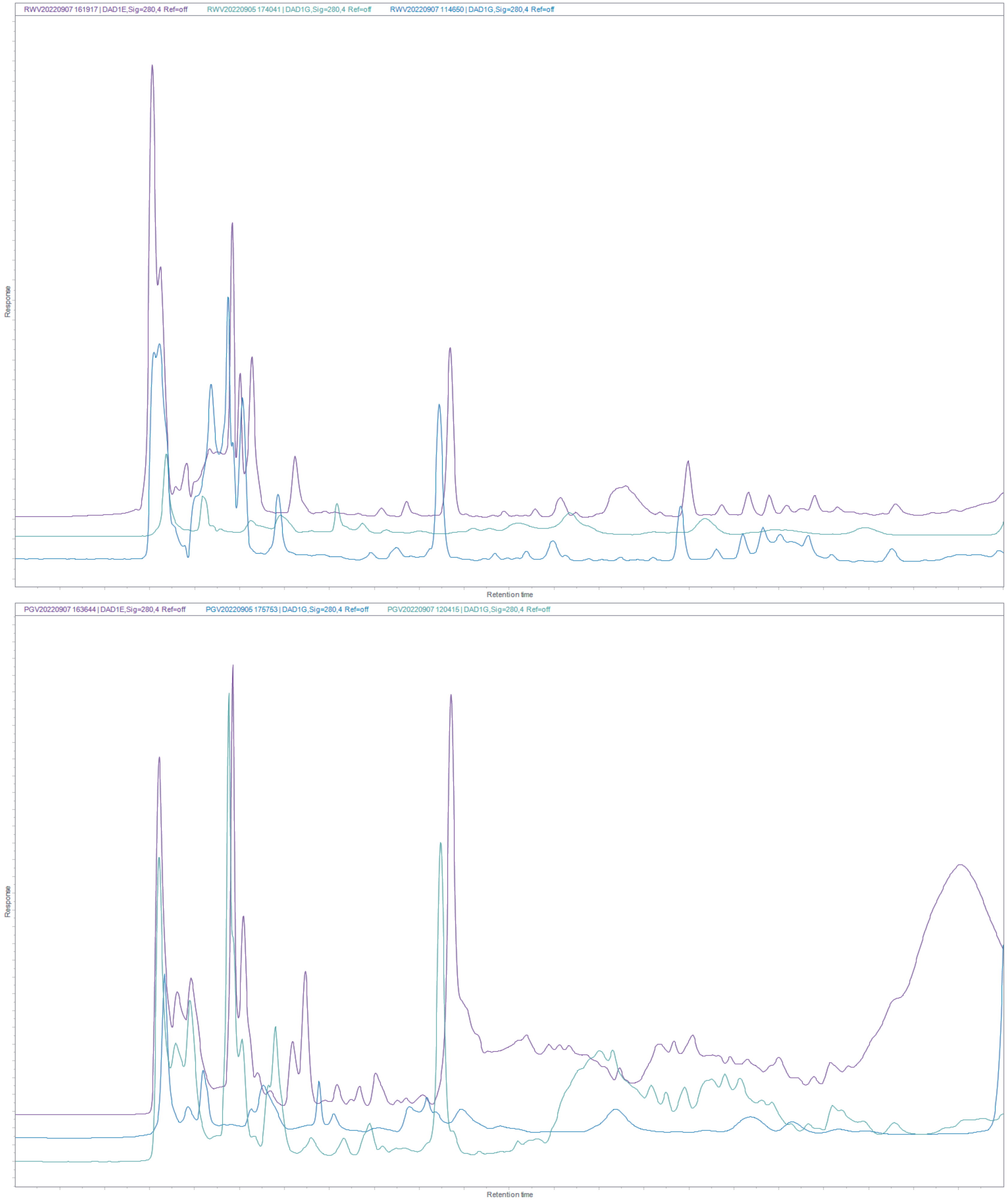


**Figure S7.** Reversed-phase HPLC chromatograms of triplicate samples of RWV2 and PV2, at 280nm

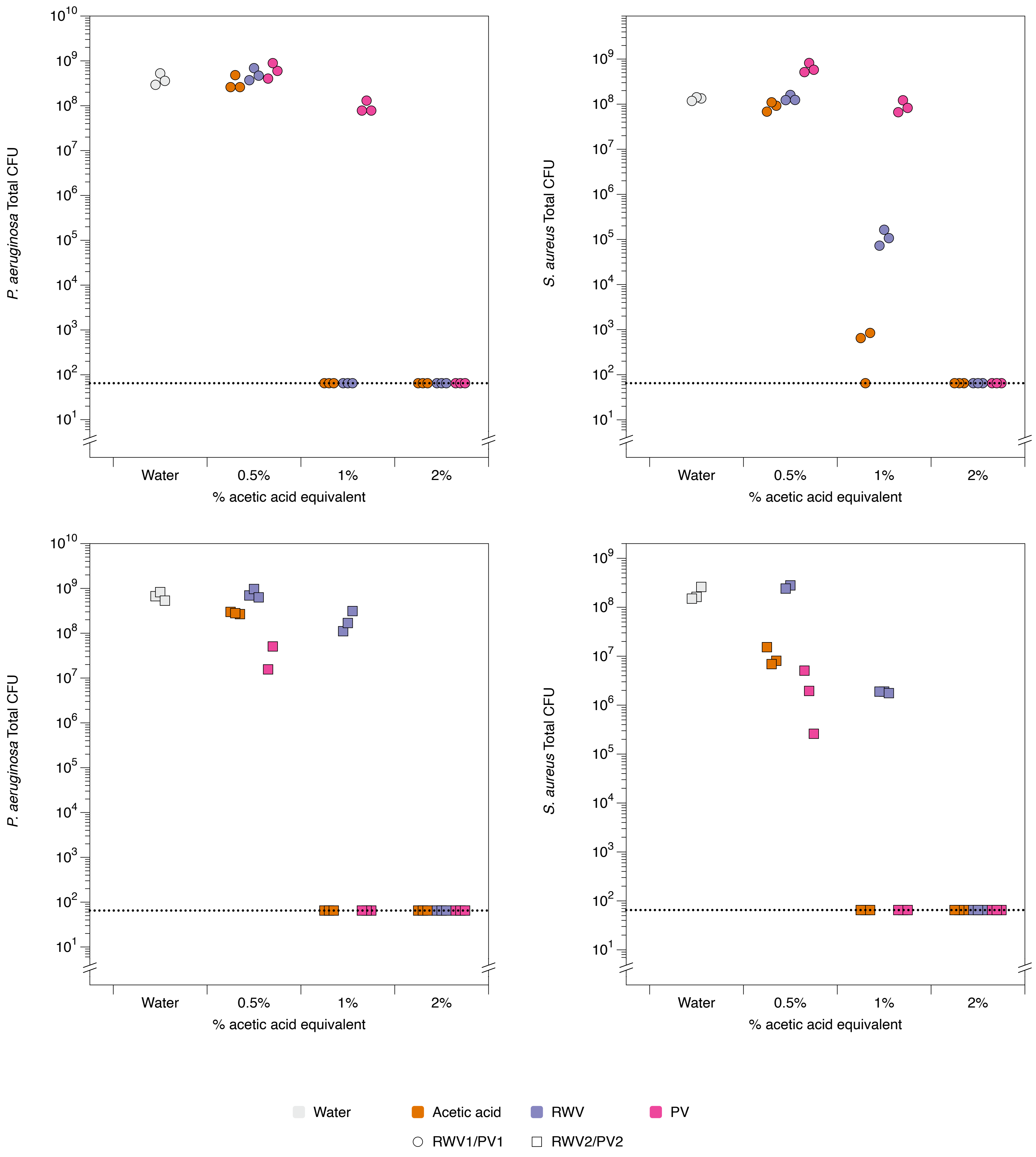


**Figure S8. Dose response of mature biofilms to acetic acid, red wine vinegars and pomegranate vinegars.** Triplicate synthetic wounds containing mature biofilms of either *P. aeruginosa* PA14 or *S. aureus* Newman were topically treated with water, acetic acid at 0.5%, 1% or 2%, or RWV1, RWV2, PV1 or PV2 at concentrations containing 0.5%, 1% or 2% acetic acid. After 24 hours' treatment, wounds were enzymatically digested to release bacteria, serially diluted and plated out to count colonies. Colony forming units (CFU) are used to estimate the number of viable bacterial cells in the biofilms. Raw data and R code are provided in the supplementary data (Document S1).
